# Coronatine contributes *Pseudomonas cannabina* pv. *alisalensis* virulence by overcoming both stomatal and apoplastic defenses in dicot and monocot plants

**DOI:** 10.1101/2020.08.19.256685

**Authors:** Nanami Sakata, Takako Ishiga, Shunsuke Masuo, Yoshiteru Hashimoto, Yasuhiro Ishiga

**Author notes:** For correspondence: Yasuhiro, Ishiga Address: Faculty of Life and Environmental Sciences, University of Tsukuba, 1-1-1, Tennodai, Tsukuba, Ibaraki 305-8572, Japan.

## Abstract

*P. cannabina* pv. *alisalensis* (*Pcal*) is a causative agent of bacterial blight of crucifer including cabbage, radish, and broccoli. Importantly, *Pcal* can infect not only a wide range of Brassicaceae, but also green manure crops such as oat. However, *Pcal* virulence mechanisms have not been investigated and are not fully understood. We focused on coronatine (COR) function, which is one of the well-known *P. syringae* pv*. tomato* DC3000 virulence factors, in *Pcal* infection processes on both dicot and monocot plants. Cabbage and oat plants dip-inoculated with a *Pcal* KB211 COR mutant (Δ*cmaA*) exhibited reduced virulence compared to *Pcal* WT. Moreover, Δ*cmaA* failed to reopen stomata on both cabbage and oat, suggesting that COR facilitates *Pcal* entry through stomata into both plants. Furthermore, cabbage and oat plants syringe-infiltrated with Δ*cmaA* also showed reduced virulence, suggesting that COR is involved in overcoming not only stomatal-based defense, but also apoplastic defense. Indeed, defense related genes, including *PR1* and *PR2*, were highly expressed in plants inoculated with Δ*cmaA* compared to *Pcal* WT, indicating that COR suppresses defense-related genes of both cabbage and oat. Additionally, SA accumulation increases after Δ*cmaA* inoculation compared to *Pcal* WT. Taken together, COR contributes to cause disease by suppressing stomatal-based defense and apoplastic defense in both dicot and monocot plants. This is the first study to investigate COR functions in the interaction of *Pcal* and different host plants (dicot and monocot plants) using genetically and biochemically defined COR deletion mutants.

**Author summary:** Disease outbreaks caused by new *Pseudomonas syringae* isolates are problems worldwide. *P. cannabina* pv. *alisalensis* (*Pcal*) causes bacterial blight on a wide range of cruciferous plants and bacterial brown spot on oat plants. Although *P. syringae* deploys a variety of virulence factors, *Pcal* virulence factors have not been investigated. We focused on coronatine (COR) function, which is one of the well-known *P. syringae* virulence factors. COR is a non-host-specific phytotoxin and contributes to *P. syringae* growth and lesion formation or expansion in several host plants. COR function has been mainly studied in the model pathogen *P. syringae* pv. *tomato* DC3000 and model the plant *Arabidopsis thaliana*. Thus, COR roles in *Pcal* infection especially on monocot plants have not been well studied. Therefore, we investigated COR role in *Pcal* interaction with both dicot and monocot plants. Here, we revealed that COR functions as a multifunctional suppressor to manage *Pcal* virulence on both plants.

## Introduction

*Pseudomonas syringae* is divided into approximately 60 pathovars which cause a of symptoms including blight, cankers, leaf spots, and galls on different plants [1]. Successful plant infection by *P. syringae* includes multiple processes which involve colonization, entry into the host plant through natural opening sites such as stomata and wounds, establishment and multiplication in the apoplast, and disease symptom production [1, 2]. In the disease cycle, *P. syringae* deploys various virulence factors for successful infection. *P. syringae* pv. *tomato* DC3000 (*Pst* DC3000) studies have revealed a large virulence factor repertoire, including the type III secretion system (TTSS) and the phytotoxin coronatine (COR) [2–5].

COR is a non-host-specific phytotoxin produced by multiple *P. syringae*. COR contributes to *P. syringae* growth and lesion formation or expansion in several host plants, including ryegrass, soybean, tomato, and several crucifers [3, 6–10]. COR is a hybrid molecule consisting of two components: coronafacic acid (CFA) and coronamic acid (CMA) [10–12]. The CFA and CMA moieties are linked through an amide bond. COR functions as a structural and functional analogue of jasmonic acid JA) and related signal compounds, such as methyl jasmonic acid (MeJA) and JA-isoleucine [13–15]. COR inhibits stomatal closure through guard-cell specific of NADPH oxidase-dependent reactive oxygen species (ROS) production triggered by bacterial microbe-associated molecular patterns (MAMPs), abscisic acid (ABA), or darkness [16–20]. The plant hormones salicylic acid (SA) and JA play major roles in defense response activation to pathogens [21–23]. SA and JA have an antagonistic relationship [22, 24], and because of the SA-JA crosstalk, COR has roles during infection, including facilitating bacterial entry through stomata, promoting bacterial multiplication and persistence *in planta*, and disease symptom induction [16, 17, 25–29].

COR biosynthesis has been well investigated in *P. syringae* pv. *glycinea* PG4180 and *Pst* DC3000 [4, 30–32]. In these strains, *corR* encodes a response regulator which positively regulates CFA and CMA biosynthesis [32, 33]. In addition to *corR*, *hrpL* encodes an alternate RNA polymerase sigma factor (σ^L^) required for TTSS expression, which also regulates COR biosynthesis [32, 34]. In *Pst* DC3000, *hrpL* regulates the COR biosynthesis cluster via *corR* [32]. These results indicate a close regulatory relationship between COR production and TTSS expression [2, 35].

*P. cannabina* pv. *alisalensis* (*Pcal*) is a causative agent of bacterial blight of crucifer including cabbage, radish, and broccoli [36–40]. *Pcal* can infect not only a wide range of Brassicaceae but also green manure crops such as oat (*Avena strigosa*) and causes brown spot on oat plants [41]. *Pcal* also produces COR [41]. Furthermore, acibenzolar-S-methyl (ASM), a well-known plant defense activator, protects cabbage and Japanese radish from *Pcal* KB211 by activating stomatal-based defense [42–44]. These results imply that overcoming stomatal-based defense is important for successful *Pcal* infection. However, studies of COR function in *Pcal* KB211 have not been conducted. Moreover, some cabbage virulence factors were also needed for oat infection, but others were not [45]. Elizabeth and Bender [8] examined the role of COR in *Pst* DC3000 virulence on different hosts including collard and turnip. COR functioned as a virulence factor in both plants, but COR was essential for *Pst* DC3000 to maintain high populations in turnip, but not in collard [8], indicating that COR may function differentially in different host plants. We investigated the role of COR in *Pcal* KB211 infection processes in different host plants, including cabbage and oat. Here, we demonstrated that COR contributes to disease by suppressing stomatal-based and apoplastic defenses on both cabbage and oat plants.

## Results

### *P. cannabina* pv. *alisalensis* KB211 causes disease on both cabbage and oat

Importantly, *Pcal* KB211 can cause disease on both cabbage and oat [41]. Firstly, we observed disease symptoms and measured bacterial populations in dip-inoculated plants. Cabbage inoculated with *Pcal* KB211 exhibited water-soaked lesions at early time points and showed chlorotic and necrotic lesions at 5 days post-inoculation (dpi) (Fig 1A). Bacterial populations in cabbage increased gradually (Fig 1B). Unlike cabbage, oat showed brown leaf spots at 5 dpi (Fig 1C), and bacterial populations suddenly increased between 3 dpi and 4 dpi (Fig 1D). These results indicate that cabbage and oat differ considerably with respect to symptom development and bacterial multiplication during *Pcal* KB211 infection.

**Figure 1.**
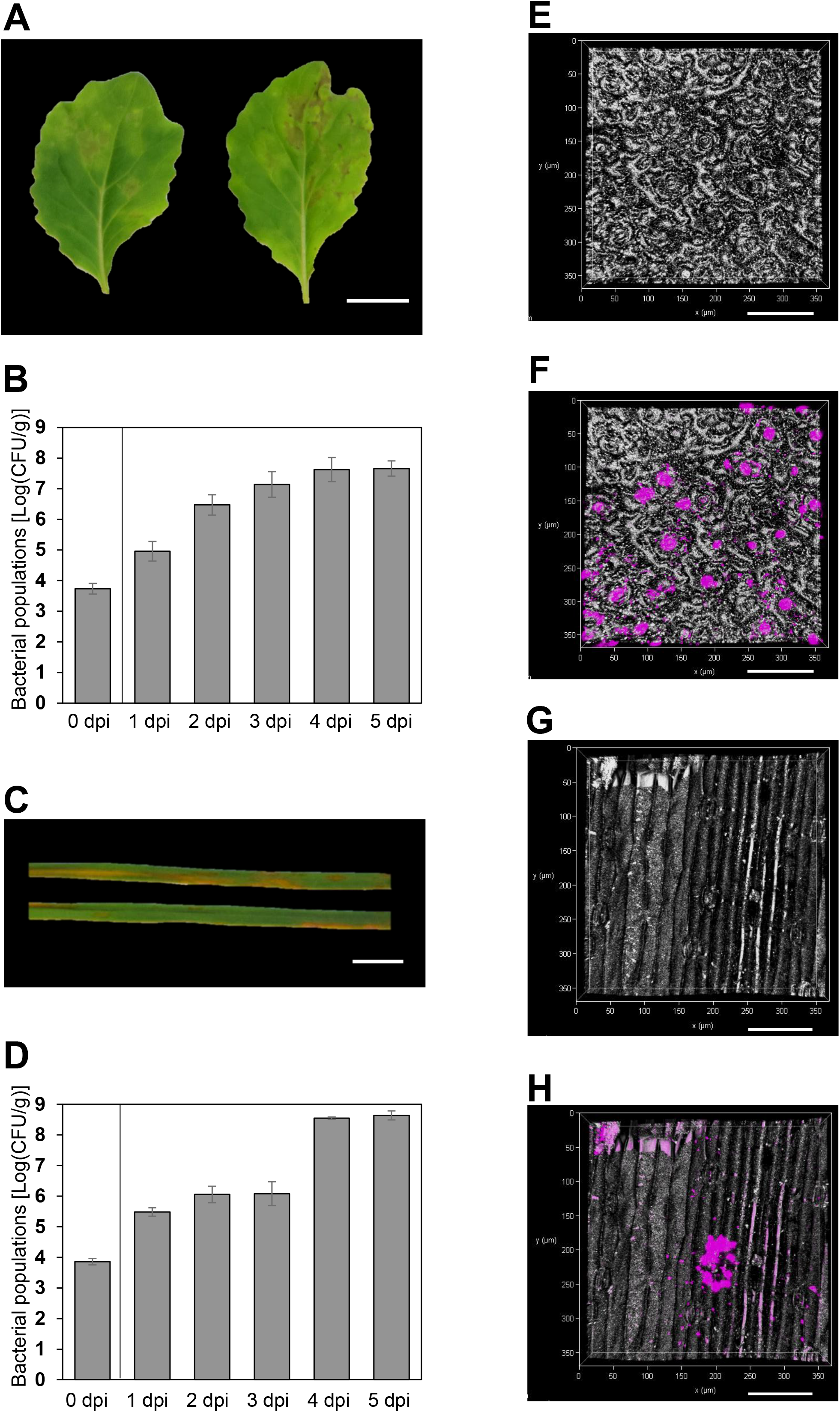
Disease phenotypes and population dynamics of *Pseudomonas cannabina* pv. *alisalnesis* KB211 in cabbage and oat. **(A) and (C)** Symptoms on cabbage (A) and oat (C) leaves dip-inoculated with *Pcal* KB211. Cabbage and oat were dip-inoculated with 5 × 10^7^ CFU/ml of inoculum. The leaves were photographed at 5 dpi. Scale bar shows 1 cm. **(B) and (D)** *Pcal* KB211 total populations on cabbage (B) and oat (C) leaves when dip-inoculated with 5 × 10^7^ CFU/ml. Vertical bars indicate the standard error for three independent experiments. **(E)-(H)** Bacterial cell location on cabbage (F) and oat (H) leaves. (E) and (G) shows the leaf surface without bacteria cells on cabbage and oat, respectively. Cabbage and oat were dip-inoculated with bacterial suspension (5 × 10^8^ CFU/ml) of tdTomato-labeled *Pcal* KB211 inoculum containing 0.025% SilwetL-77. Inoculated leaves were observed 3 dpi by using a Leica TCS SP8 confocal microscope equipped with a white light laser (Leica, Wetzlar, Germany), showing bacterial colonization of the substomatal chamber. Scale bar shows 100 μm.

We also observed the leaf surface at 3 dpi by confocal microscope, and observed multiple stomata on the cabbage leaf surface (Fig 1E). A *Pcal* KB211 bacterial colony was detected in the substomatal chamber in cabbage (Fig 1F). Conversely, oat has fewer and larger stomata compared to cabbage (Fig 1G), however bacterial colonies was also observed around the stomata (Fig 1H). These results suggest that *Pcal* KB211 enters mainly through stomata. Thus, overcoming stomatal-based defense might be critical steps in the *Pcal* KB211 infection process on cabbage and oat.

### COR biosynthesis-related genes are greatly expressed during *Pcal* KB211 infection

COR facilitates bacterial entry through stomata [16]. Therefore, we hypothesized that COR is important for *Pcal* KB211 infection processes. We investigated the expression profiles of COR biosynthesis-related genes, including *cmaA, cfl, corR*, and *hrpL,* which were greatly expressed during *Pcal* KB211 infection on both cabbage and oat compared to culture conditions (Figs 2A, 2B, 2C, and 2D). All genes were greatly expressed at 4 hpi, at which time point bacteria are supposed to overcome stomatal-based defense (Figs 2A, 2B, 2C, and 2D). Moreover, these genes also showed great expression at 48 hpi (Figs 2A, 2B, 2C, and 2D). Therefore, we hypothesized that COR is involved in overcoming both early stages and later infection processes.

**Figure 2.**
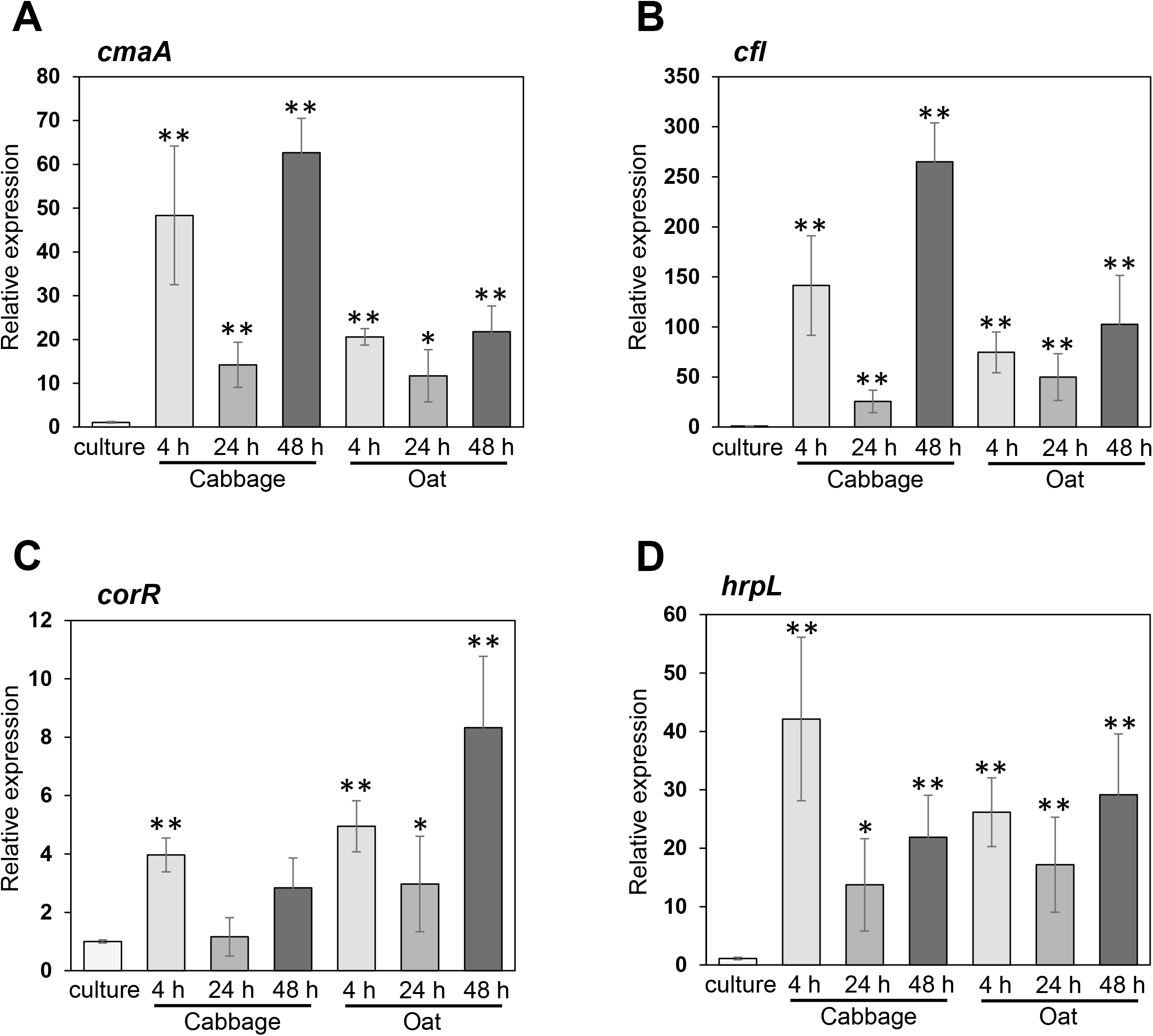
Expression profiles of *Pcal* KB211 COR biosynthesis related genes during infection on cabbage and oat plants. **(A)-(D)** Expression profiles of *cmaA* (A), *cfl* (B), *corR* (C), and *hrpL* (D) genes in liquid LB broth (culture) or in dip-inoculated cabbage and oat plants (5 × 10^5^ CFU/ml) at 4, 24, and 48 h. Total RNA was extracted for use in real-time quantitative reverse transcription-polymerase chain reaction (RT-qPCR) with gene-specific primer sets. Expression was normalized using *oprE* and *recA*. Vertical bars indicate the standard error for three biological replicates. Asterisks indicate a significant difference from culture cells in a *t* test (\**p* < 0.05, ** *p* < 0.01).

### COR contributes to disease on cabbage and oat

To assess the importance of COR in *Pcal* KB211 virulence, we constructed a COR defective *cmaA* mutant (Δ*cmaA*), and confirmed it does not produce COR by HPLC assay (S1 Fig). To investigate whether COR plays an important role in causing disease on cabbage and oat, we dip-inoculated cabbage and oat with *Pcal* KB211 wild-type (*Pcal* WT) and Δ*cmaA*. Cabbage leaves inoculated with Δ*cmaA* exhibited no necrosis and developed less chlorosis (Fig 3A). We also investigated whether COR contributes to bacterial multiplication in cabbage. Δ*cmaA* populations were significantly reduced compared to *Pcal* WT at 3 dpi and 5 dpi (Fig 3B). Δ*cmaA* also exhibited reduced disease symptoms and reduced bacterial multiplication in oat compared to *Pcal* WT at 3 dpi and 5 dpi (Figs 3C and 3D). Moreover, to further investigate whether *Pcal* WT-induced disease-associated necrotic cell death is associated with accelerated reactive oxygen species (ROS), 3,3’-diaminobenzine (DAB) staining was carried out to detect H_2_O_2_. ROS accumulation was detected in dip-inoculated cabbage and oat in response to *Pcal* WT, but not Δ*cmaA* (Figs 4A, 4D, 4G, and 4J). ROS accumulation was detected in cabbage mesophyll cell chloroplasts inoculated with *Pcal* WT (Figs 4B and 4C), but not detected in those inoculated with Δ*cmaA* (Figs 4E and 4F). Furthermore, ROS accumulation was observed in oat mesophyll cells inoculated with *Pcal* WT (Figs 4H and 4I), but not with Δ*cmaA* (Figs 4K and 4L). These results indicate that COR facilitates ROS accumulation and causes visible disease symptoms in both cabbage and oat. Taken together, COR contributes to bacterial multiplication and disease symptoms on both cabbage and oat.

**Figure 3.**
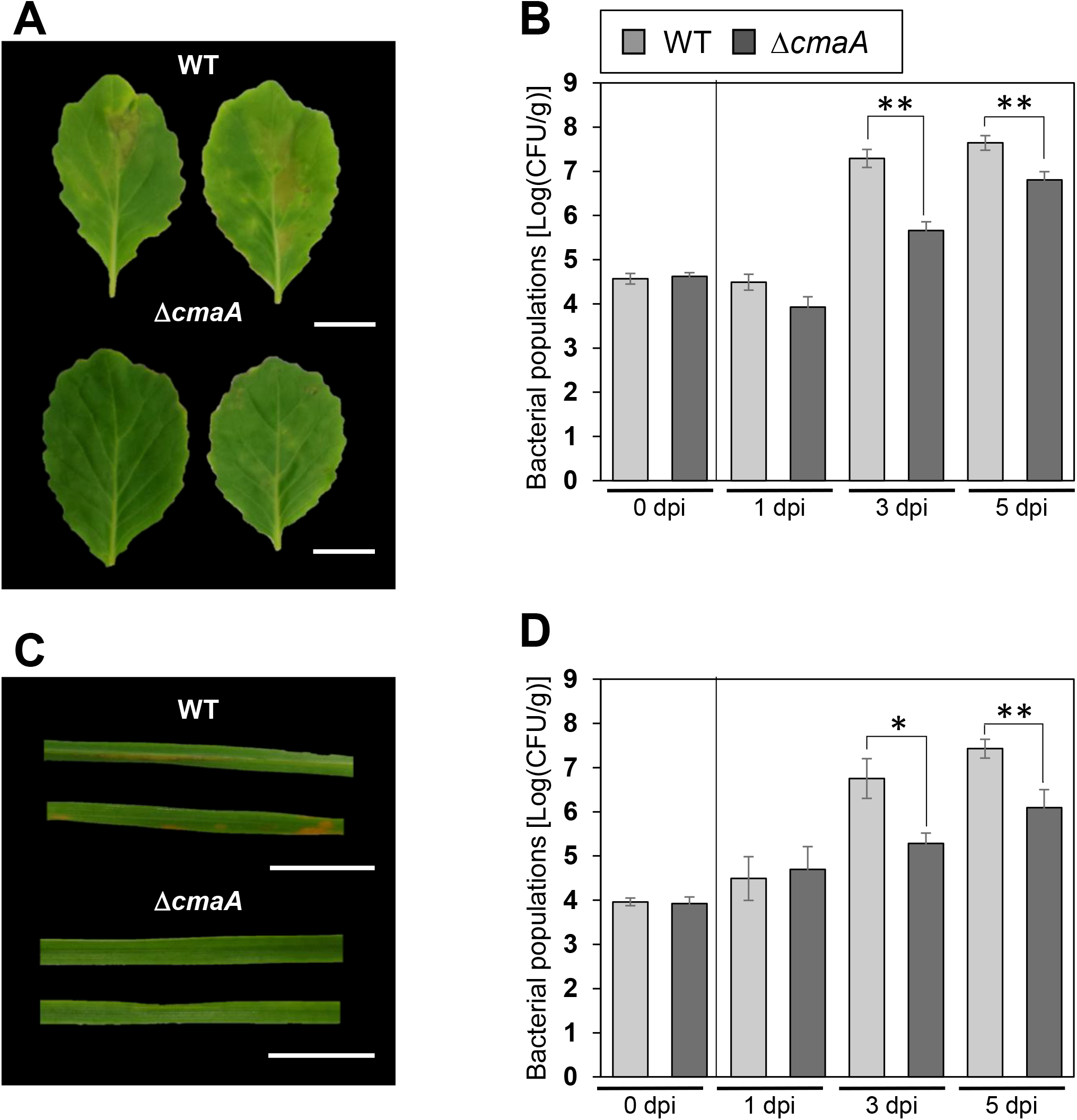
COR contributes to cause disease on cabbage and oat. **(A) and (B)** Disease symptoms (A) and bacterial populations (B) in cabbage dip-inoculated with *Pcal* WT and Δ*cmaA*. **(C) and (D)** Disease symptoms (C) and bacterial populations (D) in oat dip-inoculated with *Pcal* WT and Δ*cmaA*. Cabbage and oat were dip-inoculated with 5 × 10^7^ CFU/ml of *Pcal* KB211 containing 0.025% SilwetL-77. The leaves were photographed at 5 dpi. Vertical bars indicate the standard error for three independent experiments. Asterisks indicate a significant difference between *Pcal* WT and Δ*cmaA* in a *t* test (\**p* < 0.05, ** *p* < 0.01). Scale bar shows 1 cm.

**Figure 4.**
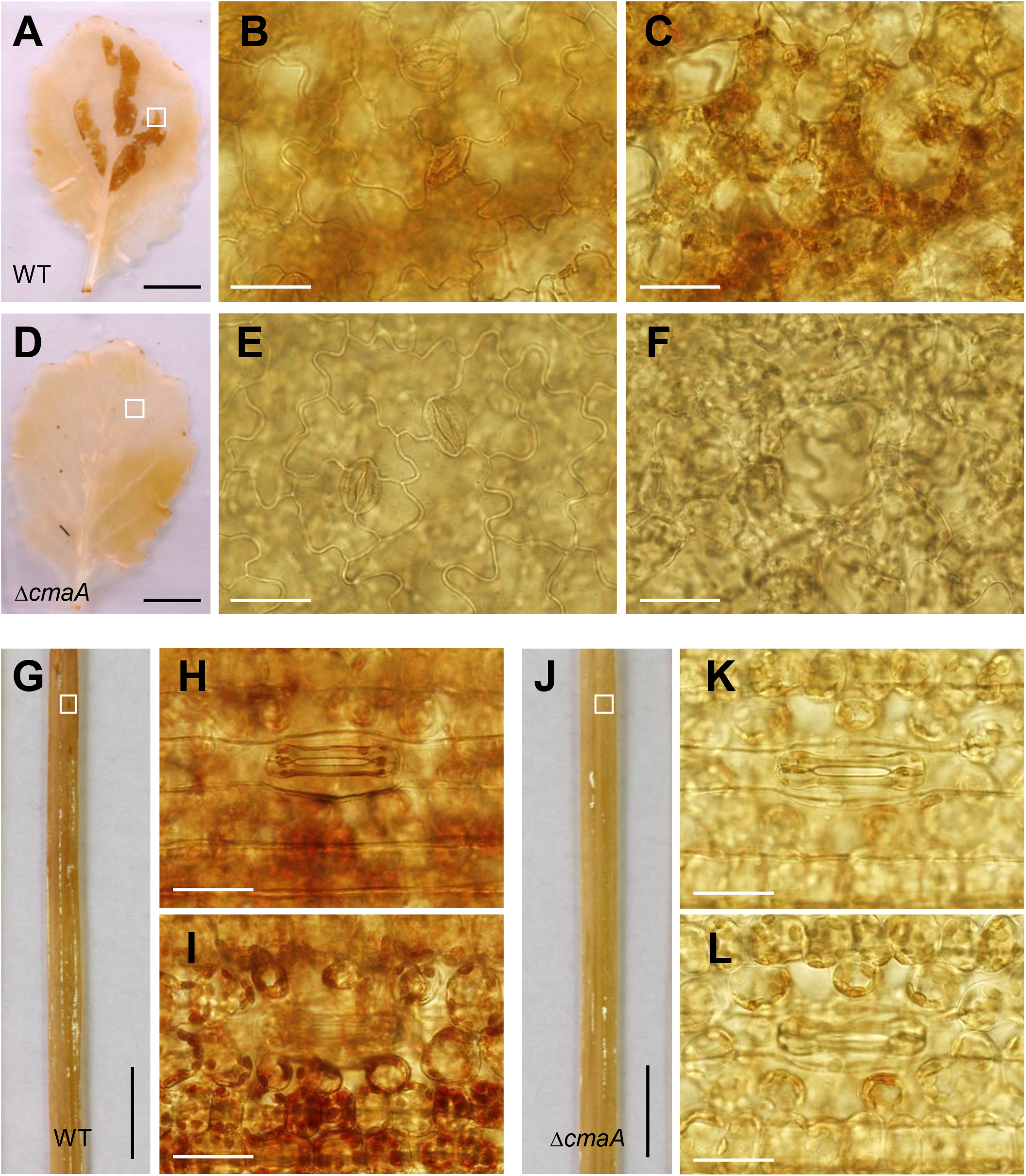
COR contributes to induce ROS accumulation. **(A)-(L)** Reactive oxygen species (ROS) accumulation visualized by staining hydrogen peroxidase using 3,3’-diaminobenzidine (DAB). DAB staining was carried out to detect H_2_O_2_ in cabbage after dip-inoculation with 5 × 10^7^ CFU/ml of *Pcal* WT (A)-(C) and Δ*cmaA* (D)-(F), and in oat dip-inoculated with 5 × 10^7^ CFU/ml of *Pcal* WT (G)-(I) and Δ*cmaA* (J)-(L). Black and white bars show 1 cm and 500 μm, respectively.

### COR reopens cabbage and oat stomata

Plants are able to respond to bacterial pathogens by actively closing the stomatal pore, so called stomatal-based defense [16, 46–51]. COR is responsible for suppressing stomatal-based defense and reopens stomata [16, 17]. Therefore, to investigate whether *Pcal* WT and Δ*cmaA* facilitate to reopen cabbage and oat stomata, we observed the stomatal aperture of both plants dip-inoculated with *Pcal* WT and Δ*cmaA* at 1 hpi and 4 hpi. Stomatal reopening was observed in both cabbage and oat inoculated with *Pcal* WT, whereas stomatal reopening was not observed in cabbage and oat inoculated with Δ*cmaA* (Figs 5A and 5B). The oat stomatal reopening rate was quite low in this experimental condition (S2 Fig). Although the stomatal response to pathogens might be different, these results suggest that COR has an important role in overcoming stomatal-based defense in *Pcal* KB211 early virulence in cabbage and oat.

**Figure 5.**
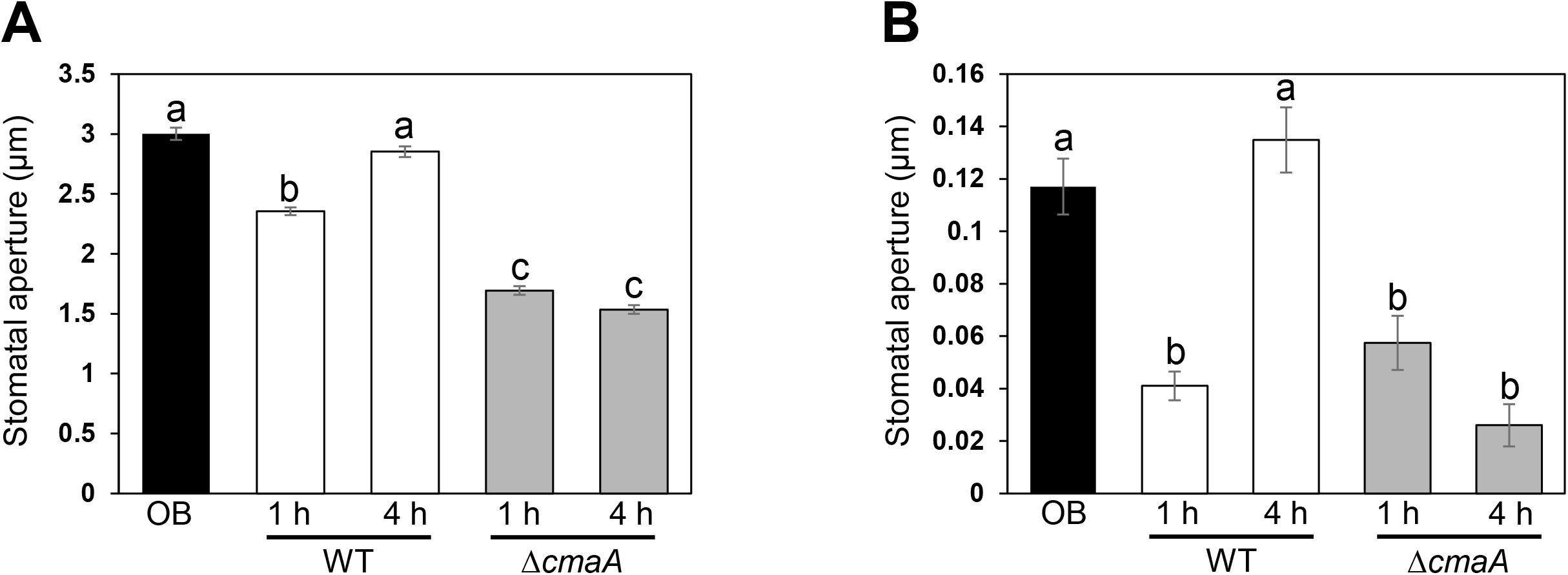
COR reopens stomata on both cabbage and oat. **(A) and (B)** Stomatal aperture width on intact cabbage (A) and oat (B) leaves 1 h and 4 h after dip-inoculation with 1 × 10^8^ CFU/ml of *Pcal* WT and Δ*cmaA*. OB indicates opening buffer control. Different letters indicate a significant difference among treatments based on a Tukey’s HSD test (*p* < 0.05).

### COR contributes to apoplastic multiplication and disease on cabbage and oat

To determine whether COR is only important for overcoming stomatal-based defense, we also conducted syringe-infiltration. Cabbage leaves inoculated with both *Pcal* WT and Δ*cmaA* showed necrosis at 5 × 10^5^ CFU/ml (S3A Fig). However, at 5 × 10^3^ CFU/ml, cabbage inoculated with Δ*cmaA* showed no symptoms, although cabbage inoculated with *Pcal* WT showed chlorosis (S3A Fig). Conversely, oat leaves inoculated with *Pcal* WT showed orange to brown spots at 5 × 10^5^ CFU/ml, and at 5 × 104 CFU/ml (S3A Fig). Oat inoculated with Δ*cmaA* showed no symptoms at all bacterial inoculum levels (S3B Fig). Therefore, we next examined bacterial multiplication in cabbage and oat inoculated with 5 × 10^3^ CFU/ml and 5 × 10^5^ CFU/ml, respectively. In cabbage, the Δ*cmaA* bacterial populations were significantly reduced at 1 and 2 dpi (Fig 6B). Interestingly, although Δ*cmaA* showed no symptoms, the bacteria multiplied almost to the same level as *Pcal* WT at 3 dpi (Figs 6A and 6B). In contrast, Δ*cmaA* showed no symptoms and did not grow until 5 dpi in oat (Figs 6C and 6D). These results indicate COR plays an important role in overcoming apoplastic defenses, and may function differentially in cabbage and oat.

**Figure 6.**
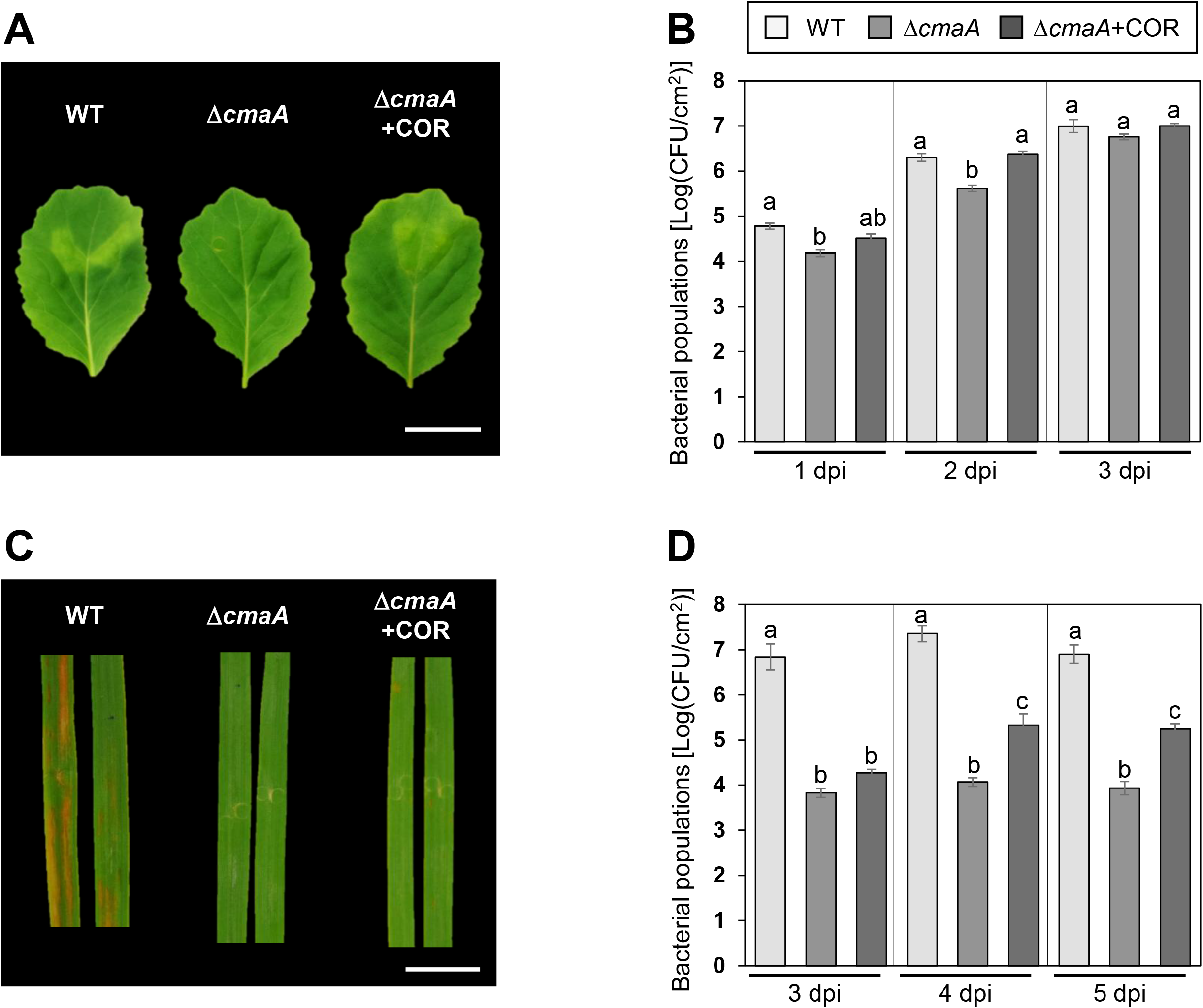
COR contributes to cause disease on cabbage and oat. **(A) and (B)** Disease symptoms (A) and bacterial populations at 1, 2, and 3 dpi (B) in cabbage syringe-inoculated with *Pcal* WT and Δ*cmaA*. Cabbage were syringe-inoculated with 5 × 10^3^ CFU/ml of *Pcal* WT and Δ*cmaA*. **(C) and (D)** Disease symptoms (C) and bacterial populations at 1, 2, and 3 dpi (D) in oat syringe-inoculated with *Pcal* WT and Δ*cmaA*. Oat were syringe-inoculated with 5 × 10^5^ CFU/ml of *Pcal* WT and Δ*cmaA*. Additionally, COR (500 μM) was co-inoculated with Δ*cmaA*. The leaves were photographed at 5 dpi. Vertical bars indicate the standard error for three independent experiments. Different letters indicate a significant difference among treatments based on a Tukey’s HSD test (*p* < 0.05). Scale bar shows 2 cm.

We further tested whether exogenous COR application restores disease susceptibility to cabbage and oat. Co-inoculation of Δ*cmaA* with COR 500 μM increased bacterial growth to a similar level as *Pcal* WT in cabbage at 2 dpi (Fig 6B). In oat, this reduced multiplication could be partly recovered by COR application from 4 dpi (Fig 6D). Therefore, these results suggest that COR is strongly involved in multiplication in oat, especially in later infection stages, compared to cabbage.

### COR regulates defense-related gene expression in cabbage and oat

To evaluate the effect of COR on defense-related gene expression in cabbage and oat, we first investigated *PR* gene expression profiles, including *PR1* and *PR2*, in response to the SA analog plant activator, acibenzolar-S-methyl (ASM). *BoPR1* and *BoPR2* expression were induced by ASM treatment in cabbage (Figs 7A, and 7B), and *AsPR1a*, *AsPR1b*, and *AsPR2* were induced in oat (Figs 7C, 7D, and 7E). Therefore, we investigated the expression profiles of these in response to *Pcal* WT and Δ*cmaA*. Cabbage *BoPR2* genes showed significantly greater expression levels at 24 h and 48 h in response to Δ*cmaA* compared to *Pcal* WT (Fig 7G). *BoPR1* genes showed no significant difference but tended to increase at 48 h when we inoculated with Δ*cmaA* compared to *Pcal* WT (Fig 7F). In oat, all *AsPR1a, AsPR1b*, and *AsPR2* genes showed significantly greater expression levels at 24 h and 48 h in response to Δ*cmaA* compared to *Pcal* WT (Figs 7H, 7I, and 7J). These results indicate that COR suppresses SA-mediated signaling pathways leading to PR protein accumulation.

**Figure 7.**
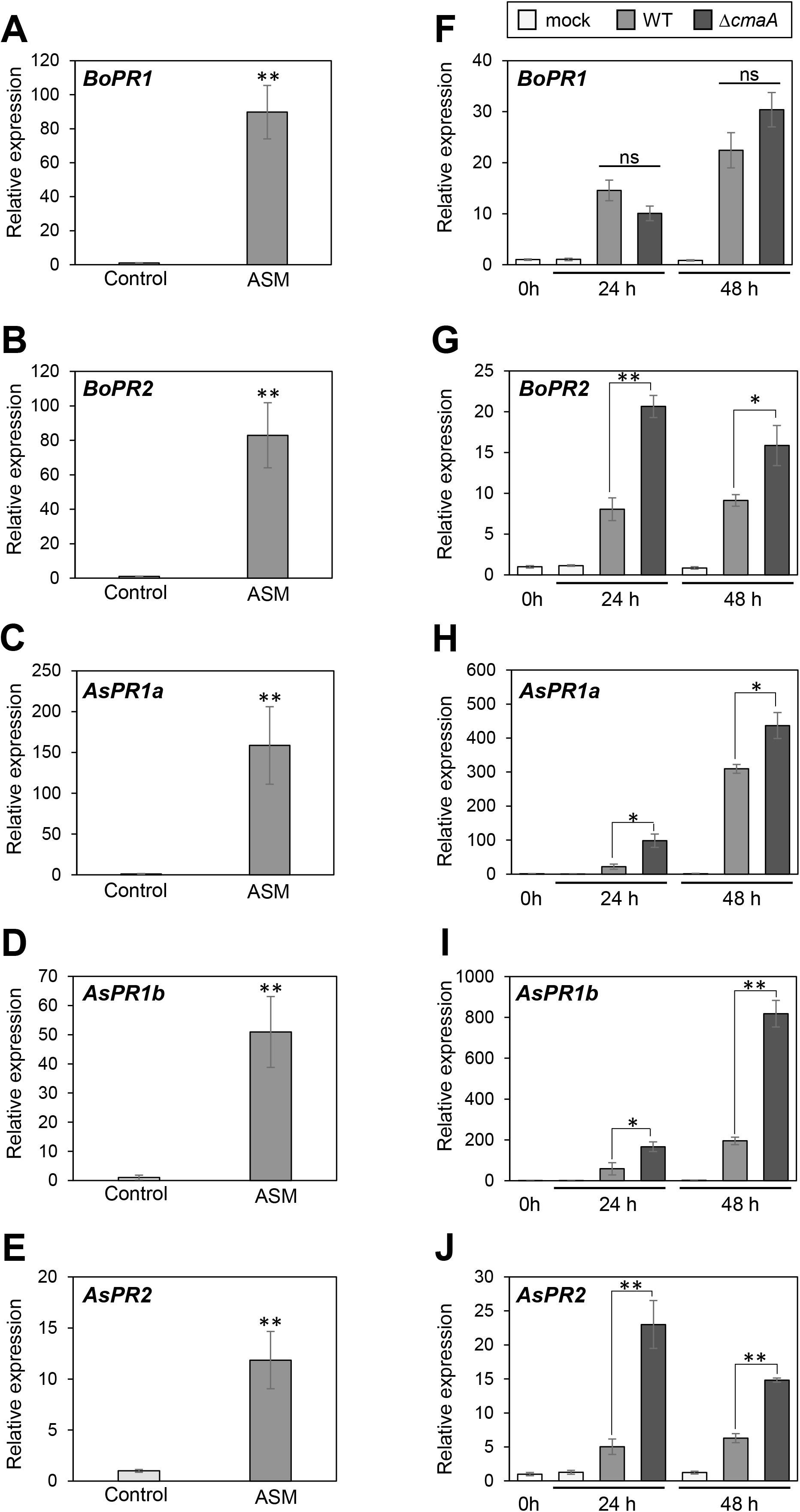
COR suppresses defense-related gene expression in cabbage and oat. **(A)-(E)** Gene expression profiles involved in cabbage and oat plant defense after ASM treatment. Cabbage and oat were treated with water as a control and ASM (100 ppm). Expression levels of *BoPR1* (A), *BoPR2* (B), *AsPR1a* (C), *AsPR1b* (D), and *AsPR2* (E) were determined using real-time quantitative reverse transcription-polymerase chain reaction (RT-qPCR) with gene-specific primer sets. Expression was normalized using *BoUBQ1* and *PsAct* in cabbage and oat, respectively. Vertical bars indicate the standard error for three biological replicates. Asterisks indicate a significant difference from the water control in a *t*-test (* *p* < 0.05; ** *p* < 0.01). ns stands for no significant difference between samples. **(F)-(J)** Gene expression profiles involved in cabbage and oat plant defense after syringe-inoculation with water (mock), *Pcal* WT, and Δ*cmaA*. Expression profiles of *BoPR1* (F), *BoPR2* (G), *AsPR1a* (H), *AsPR1b* (I), and *AsPR2* (J) were determined 24 h and 48 h after inoculation with 5 × 10^5^ CFU/ml of *Pcal* WT and Δ*cmaA*, or mock water-inoculated control using RT-qPCR with gene-specific primer sets. Expression was normalized using *BoUBQ1* and *PsAct* in cabbage and oat, respectively. Vertical bars indicate the standard error for three biological replicates. Asterisks indicate a significant difference between *Pcal* WT and Δ*cmaA* in a *t*-test (* *p* < 0.05; ** *p* < 0.01).

### COR is involved in suppression of SA biosynthesis

We demonstrated that SA-induced gene markers, including *PR1* and *PR2*, are greatly expressed in plants after inoculation with Δ*cmaA* compared to *Pcal* WT (Figs 7F, 7G, 7H, 7I, and 7J). Thus, we next examined SA accumulation levels in cabbage and oat when inoculated with *Pcal* WT and Δ*cmaA* by using LC-MS/MS. In cabbage, SA accumulation with mock treatment was less than 20 ng/g, and increased around twenty-fold at 24 h after WT inoculation (Fig 8A). Δ*cmaA* inoculation induced greater SA levels than *Pcal* WT inoculation at 24 hpi (Fig 8A), indicating that COR suppresses SA biosynthesis in cabbage within 24 h after inoculation. Conversely, SA accumulation triggered by bacterial pathogens caused subtle changes, and showed no statistical difference between *Pcal* WT and Δ*cmaA* inoculations in oat (Fig 8B). However, at 48 h, SA accumulation also tended to be suppressed by COR in oat (Fig 8B). Therefore, although COR contributes to SA suppression differently, COR can suppress SA accumulation and allow *Pcal* KB211 to multiply to great levels in both cabbage and oat.

**Figure 8.**
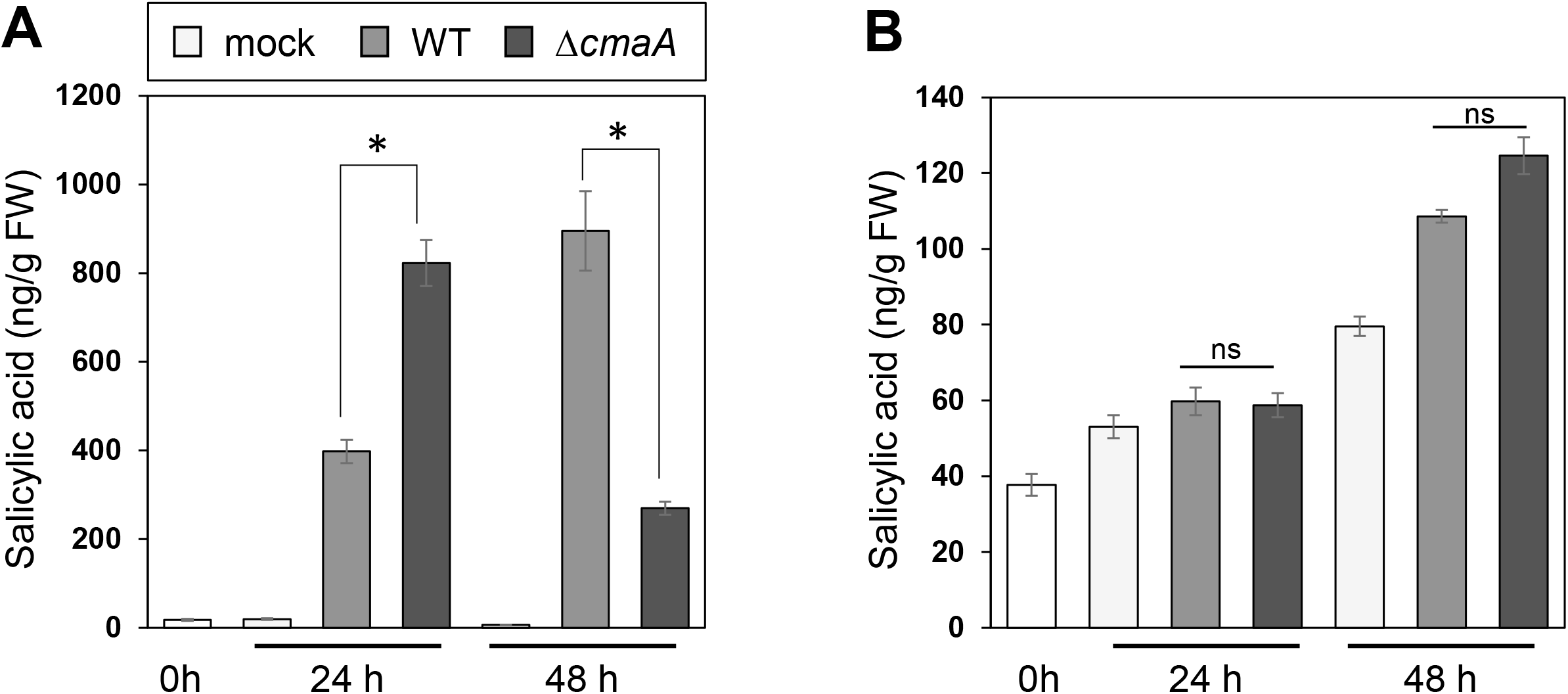
COR suppresses SA accumulation in cabbage and oat. **(A)-(B)** Total SA production in cabbage (A) and oat (B) after syringe-inoculation with 5 × 10^5^ CFU/ml of *Pcal* WT and Δ*cmaA*, or treated with water as a control. Vertical bars indicate the standard error for three biological replicates. Asterisks indicate a significant difference between plants inoculated with *Pcal* WT and Δ*cmaA* in a *t*-test (* *p* < 0.05). ns indicate no significant difference between samples.

## Discussion

Plant pathogens deploy a variety of virulence factors. *Pcal* KB211 multiplicated mainly around stomata (Figs 1), and COR biosynthesis-related genes were greatly expressed during early and later infection processes (Figs 2). Therefore, we hypothesized that COR contributes to *Pcal* KB211 virulence for overcoming both stomatal-based and apoplastic defenses. COR is involved in suppressing stomatal closure (Figs 5) and SA-mediated defenses (Figs 7 and 8) in both cabbage and oat. These results shed light on spatiotemporal COR function in different hosts, monocot and dicot.

The Δ*cmaA* multiplication defect was observed at 3 dpi, and this defect became greater at 5 dpi in both cabbage and oat (Figs 3). Δ*cmaA* was defective in reopening stomata in both cabbage and oat (Figs 5), indicating that COR suppresses stomatal-based defense and facilitates *Pcal* entry through stomata into both cabbage and oat. Melotto et al. [16] demonstrated that *Pst* DC3118 *cor* mutant populations were greatly reduced compared with those of *Pst* DC3000 when dip-inoculated on *A. thaliana*. This multiplication defect was observed at early infection stages (3 dpi), suggesting that the COR-mediated suppression of stomatal defense is critical for *Pst* DC3000 host plant infection [16]. COR is required for *Pst* DC3000 growth and persistence within tomato and *A. thaliana* [27, 31]. These results highly support our results that COR is important for disease symptom development and growth in both cabbage and oat. Elizabeth and Bender [8] investigated the role of COR in the interaction of *Pst* DC3000 and different brassicas, collard and turnip. They demonstrated that COR contribution to bacterial multiplication in collard was more subtle than in turnip [8]. However, the symptoms on collard appeared earlier than on turnip, and bacterial growth in collard reached greater levels earlier than in turnip. They speculated that COR might prolong stomatal aperture opening in turnip, but not in collard. The stomatal opening rate is quite decreased in oat in our experimental conditions (S2 Fig). Roles of stomata in plant disease resistance are not fully understood, especially in monocot plants. Zhang et al. [52] demonstrated that open stomata in rice lead to leaf blight bacterial resistance against *Xanthomonas oryzae* pv. *oryzae*, and this resistance was conferred through modulation of host water status. It is possible that stomatal-based defense response differs between cabbage and oat. Further investigations on stomatal response will be needed to understand stomatal-based defense, especially in monocot plants.

Δ*cmaA* populations were significantly less than those of *Pcal* WT in both syringe-inoculated cabbage and oat (Figs 6). These results indicate COR functions not only in suppressing stomatal-based defense but also in overcoming apoplastic defense. This multiplication defect was also observed in *A. thaliana* plants vacuum-infiltrated with the *Pst* AK7E2 (*cmaA* mutant) and *Pst* DB4G3 (*cfa6* mutant) at 4 dpi, but not at 2 dpi [31], indicating that COR may be important for apoplastic bacterial persistence. However, Melotto et al. [16] observed no multiplication defect at 3 dpi when the *cor* mutant *Pst* DC3118 was infiltrated directly into the apoplast compared to *Pst* DC3000. Additionally, Mittal and Davis [7] also showed multiplication impairment of *Pst* DC3661, a Tn*5* Cor^-^ derivative mutant, when dip-inoculated, but multiplication impairment was not observed in *A. thaliana* and tomato inoculated by infiltration. In these studies, the inoculum concentration was 1 × 10^6^ CFU/ml. These results highly motivated us to investigate the bacterial multiplication at different time points and different inoculum concentrations. Δ*cmaA*-inoculated cabbage leaves exhibited similar disease symptoms to *Pcal* WT-inoculated leaves at 5 × 10^5^ CFU/ml (S3A Fig). Conversely, oat infiltrated with Δ*cmaA* did not show any symptoms even at the greatest concentration we tested, 5 × 10^5^ CFU/ml (S3B Fig). Helmann et al. [53] demonstrated that disease symptoms were observed only when local internal population sizes in excess of about 10^5^ cells/cm^2^ were present, suggesting that bacterial concentrations above a specific “threshold” were required for visible lesion formation. Thus, the threshold might be different between cabbage and oat. Therefore, our results support the importance of selecting the proper inoculum levels depending on the host plant and bacterial pathogen.

We demonstrated that defense-related genes, including *PR1* and *PR2,* were greatly expressed when we inoculated cabbage and oat with Δ*cmaA* compared to *Pcal* WT (Figs 7), indicating that COR suppresses defense-related gene expression. Also, we showed that SA accumulation was greater in cabbage and oat inoculated with Δ*cmaA* compared to *Pcal* WT at 24 h (Fig 8A). SA accumulation in oat showed subtle changes against the bacterial pathogen, and no statistical change between the *Pcal* WT and Δ*cmaA* inoculations (Fig 8B). However, concerning bacterial multiplication, the Δ*cmaA* population was 10-fold less than those of *Pcal* WT. Therefore, taken together, our results showed that COR suppressed SA-mediated defense in both cabbage and oat. Several studies have demonstrated that COR functions to suppress SA-mediated defenses via COI1 activation, thus allowing bacteria to grow to greater densities *in planta* in dicot plants, such as *A. thaliana* and tomato [27, 29, 54, 55]. However, COR function in SA-mediated defense in monocot plants is less studied. Moreover, in monocot plants, the role of SA biosynthesis in disease resistance is still poorly understood [56]. Consistent with our results on oat, rice plants accumulated increased basal SA levels (8-37 mg/g FW) that did not change significantly upon pathogen attack [57]. In contrast, in tobacco and *A. thaliana*, basal SA levels were less (less than 100ng/g FW), but increased by two orders of magnitude following pathogen infection [58]. However, several reports indicated that SA-JA antagonism is conserved in monocot plants such as rice and wheat. In wounded rice, JA levels elevated whereas SA levels decreased, indicating negative crosstalk between the JA and SA pathways during early wounding response in rice [59]. Furthermore, Ding et al. [60] also demonstrated that as in dicot plants, SA and JA acted antagonistically in pathogen-inoculated wheat plants. Also, Iwai et al. [61] demonstrated that SA accumulation in rice was induced after probenazole (PBZ) treatment (one of the synthetic SA analogs) in adult leaves, but not in young leaves. Therefore, SA-mediated defense might be quite different depending on leaf age and plant species. To understand the mechanism by which COR suppresses SA accumulation, further investigation of SA-JA interactions in monocots will be necessary.

Suppression of SA-mediated defense by COR results in reduced apoplastic defense in cabbage and oat (Figs 6, 7, and 8). However, COR may contribute to suppress not only SA-mediated defense, but also other apoplastic defenses, such as plant secondary metabolites synthesis. Geng et al. [62] showed that COR inhibited the SA-independent pathway, contributing to callose deposition by reducing accumulation of an indole glucosinolate. Consistent with this study, when *A. thaliana* was challenged by *Pst* DC3000, COR affected gene expression involved in the synthesis of Trp-derived glucosinolates [63]. Brassica species, including cabbage, produce elevated levels of secondary metabolites all proposed to originate from indole glucosinolate [64]. Therefore, there is a possibility that COR regulates secondary metabolites in cabbage. These plant-derived secondary metabolites are known to have an important role in plant defense through their antimicrobial activities. Indeed, brassinin, one of the indole-sulfur phytoalexins produced in cabbage, can cause 50% growth inhibition of *Lectosphaeria maculans* [65]. Moreover, Wang et al. [66] reported that an *A. thaliana* secondary metabolite, sulforaphane, directly targets TTSS expression to inhibit bacterial virulence. These studies suggest that plant secondary metabolites could have a major role in apoplastic defense. In oat, a group of phenolic antioxidants termed avenathramides and saponins, have been well characterized as phytoalexins [67–71]. Avenantheramide levels in vegetation were enhanced by BTH treatment, which is a SA analog same as ASM [68]. This study implied the possibility that COR might suppress avenanthramide biosynthesis, because of the SA-JA antagonistic relationship. Therefore, these is a possibility that COR contributes to overcome apoplastic defense not only by suppressing SA-mediated defense, but also by modulating plant secondary metabolites. Further exploration will be necessary to better understand the potential function of COR in apoplastic defense.

In summary, COR contributes to *Pcal* KB211 virulence on dicot and monocot plants in both early and later stages of the infection process by suppressing stomatal-based and apoplastic defenses. Moreover, we suggest that COR might contribute differentially to *Pcal* KB211 virulence between monocot and dicot plants. This differential might be strongly involved in apoplastic defenses in each host plant. We need to investigate the possibilities which lead to these differential results to understand the potential function of COR and host optimization.

## Materials and methods

### Bacterial strains, plasmids, and growth conditions

The bacterial strains and plasmids used in this study are described in S1 Table. *Pseudomonas cannabina* pv. *alisalensis* strain KB211 (*Pcal* KB211) was used as the pathogenic strain to inoculate cabbage and oat. *Pcal* KB211 wild-type (WT) and the Δ*cmaA* mutant were grown on King’s B (KB; [72]) medium at 28°C, and *E. coli* strains were grown on Luria-Bertani (LB; [73]) medium at 37°C. Antibiotics used for selection of *Pcal* and *E. coli* strains included (in μl/ml): rifampicin, 50; kanamycin, 25; ampicillin, 100; chloramphenicol, 10. Before *Pcal* plant inoculation, bacteria were suspended in sterile distilled H_2_O, and the bacterial cell densities at 600 nm (OD_600_) were measured using a Biowave CO8000 Cell Density Meter (Funakoshi, Tokyo, Japan).

### Generation of tdTomato-labeled Pcal KB211 and microscopic observation

A pDSK-*tdTomato* plasmid was generated by replacing the *GFPuv* gene with *tdTomato* (S1 Table) in pDSK-*GFPuv* [74] for tdTomato fluorescent protein expression. The pDSK-*tdTomato* vector was introduced into *Pcal* KB211 by electroporation. Dip-inoculated cabbage and oat leaves were directly imaged using a Leica TCS SP8 confocal microscope equipped with a white light laser (Leica, Wetzlar, Germany). A reflected image of the leaf surface was obtained by illuminating the sample with 480 nm wavelength, and reflected light was detected through 475-486 nm laser lines. tdTomato red fluorescence was detected by 554-611 nm laser lines.

### Plant materials

Cabbage (*Brassica oleracea* var. *capitate*) cv. Kinkei 201 and oat (*Avena strigosa*) cv. Hayoat plants were used for *Pcal* virulence assays. Cabbage and oat were grown from seed at 23-25°C with a light intensity of 200 μEm^−2^s^-1^ and a 16 h light/8 h dark photoperiod. Cabbage and oat plants were used for dip-inoculation assays around two weeks after germination and for syringe-inoculation assays around three weeks after germination.

### Generation of Δ*cmaA* mutant

The mutants were generated as described previously [75]. The genetic regions containing *cmaA* and surrounding regions were amplified using PCR primer sets (S2 Table) that were designed based on the *Pcal* ES4326 registered sequence with Prime Star HS DNA polymerase (TaKaRa, Otsu, Japan), then dA was added to the 3’ end of the PCR product with 10 x A-attachment mix (TOYOBO, Osaka, Japan). The resultant DNA was inserted into the pGEM-T Easy vector (Promega, Madison, WI, USA). The recombinant plasmid DNA was then used to obtain pGEM-*cmaA* as templates, and inverse PCR was carried out using a primer set (S1 Table) to delete an open reading frame for *cmaA*. Then, the PCR product and template DNA were digested with BamHI and DpnI. The resultant DNA was self-ligated with T4 DNA ligase (Ligation-Convenience kit, Nippon Gene, Tokyo, Japan). The *cmaA*-deleted DNA constructs were introduced into the EcoRI site of the mobilizable cloning vector pK18*mobsacB* [76]. The resulting plasmids containing the DNA fragment lacking *cmaA* were then transformed into *E. coli* S17-1. The deletion mutant was obtained by conjugation and homologous recombination according to the method previously reported [77]. Transconjugants were selected on KB agar containing 30 μg/ml of kanamycin (Km) and 30 μg/ml of rifampicin (Rif). We generated mutants by incubating the transconjugants on a KB agar plate containing 25 μg/ml of rifampicin and 10% sucrose. The specific deletions in the Δ*cmaA* mutants were confirmed by PCR using primers (S1 Table).

### Bacterial inoculation methods

To assay for disease on cabbage and oat seedlings, dip-inoculation was conducted by soaking seedlings in bacterial suspensions (5 × 10^7^ CFU/ml) containing 0.025% Silwet L-77 (OSI Specialities, Danbury, CT, USA). The seedlings were then incubated in growth chambers at 85-95% RH for the first 24 h, then at 80-85% RH for the rest of the experimental period. Disease symptom pictures on cabbage and oat seedlings were taken at 5 days post-inoculation. For syringe-inoculation, bacteria were suspended at a final concentration ranging from 5 × 10^2^ to 5 × 10^5^ CFU/ml, and infiltrated with a 1-ml blunt syringe into leaves. The plants were then incubated at 70-80% RH for the rest of the experimental period. Leaves were removed and photographed at 5 days post inoculation.

To assess bacterial growth in cabbage and oat seedlings, the internal bacterial population was measured after dip-inoculation. Inoculated seedlings were collected, and a total of two inoculated leaves were measured chronologically on both plants. The leaves were surface-sterilized with 10% H_2_O_2_ for 3 minutes. After washing three times with sterile distilled water, the leaves were homogenized in sterile distilled water, and diluted samples were plated onto solid KB agar medium. For syringe-inoculation, to assess bacterial growth in cabbage seedlings, the internal bacterial population was measured after syringe-inoculation. Leaf discs were harvested using a 3.5 mm-diameter cork-borer from syringe-infiltrated leaf zones. To assess bacterial growth in oat, leaf pieces were cut from syringe-infiltrated leaf zones and the area (cm^2^) measured. Leaf extracts were homogenized in sterile distilled water, and diluted samples were plated onto solid KB agar medium. Two or three days after dilution sample plating, the bacterial colony forming units (CFUs) were counted and normalized as CFU per gram or CFU per cm^2^, using the total leaf weight or leaf square meters. The bacterial populations at 0 dpi were estimated using leaves harvested 1 hpi without surface-sterilization. The bacterial populations were evaluated in at least three independent experiments.

### Real-time quantitative RT-PCR

For expression profiles of cabbage defense genes in response to ASM, cabbage and oat were treated by drenching the soil with ASM. After 24 h, total RNA was extracted from leaves and purified using RNAiso Plus (TaKaRa) according to the manufacture’s protocol. For expression profiles of *Pcal* KB211 genes in culture, bacteria were grown in LB broth for 24 h, adjusted to an OD_600_ of 0.1, and grown again in LB broth for 3 hours. Then, bacteria were transferred to fresh mannitol-glutamate broth (MG; [78]) and grown for 15 min. Bacterial RNA was extracted using ReliaPrep RNA Cell Miniprep System Kit (Promega, WI, USA) according to the manufacture’s protocol. To analyze *Pcal* KB211 gene expression profiles during infection, we dip-inoculated cabbage and oat with *Pcal* KB211 at 1 x 10^8^ CFU/ml for 4, 24, and 48 h, then the total RNAs including plant and bacterial RNAs were extracted from infected leaves and purified using RNAiso Plus (TaKaRa) according to the manufacture’s protocol. The real-time quantitative RT-PCR (RT-qPCR) were done as described previously [79]. Two micrograms of total RNA were treated with gDNA Remover (TOYOBO, Osaka, Japan) to eliminate genomic DNA, and the DNase-treated RNA was reverse transcribed using the ReverTra Ace qPCR RT Master Mix (TOYOBO). The cDNA (1:10) was then used for RT-qPCR using the primers shown in Supplementary Table S1 with THUNDERBIRD SYBR qPCR Mix (TOYOBO) on a Thermal Cycler Dice Real Time System (TaKaRa). *Pcal* KB211 *oprE* and *recA* were used to normalize gene expression. Cabbage *UBIQUITIN EXTENSION PROTEIN 1* (*BoUBQ1*) and Oat *Actin* (*PsAct*) were used as internal controls to normalize gene expression in cabbage and oat, respectively.

### Hydrogen peroxidase detection

Hydrogen peroxidase generation was detected by 3,3'-diaminobenzidine (DAB) staining as described previously [28, 80]. After inoculation with *Pcal* WT and Δ*cmaA,* the leaves were placed in DAB-HCL (pH 3.8) at 1 mg/ml. After incubation for 6 h at room temperature, chlorophyll was removed with 95% ethanol.

### Stomatal assay

A modified method was used to assess stomatal response as described previously [81]. Briefly, cabbage and oat were grown for around 2 weeks after germination as described previously. *Pcal* WT and Δ*cmaA* were grown at 28°C for 24 h on KB agar, then suspended in distilled water to an OD_600_ of 0.2 (1 × 10^8^ CFU/ml). Dip-inoculated cabbage and oat leaves were directly imaged at 1 hpi and 4 hpi using a Nikon optical microscope (Eclipse 80i). The aperture width of at least 300 and 600 stomata was measured on cabbage and oat, respectively. The average and standard error for the stomatal aperture width were calculated. The stomatal apertures were evaluated in at least three independent experiments.

### SA quantification by RP-LC-ESI-MS/MS

*Pcal* WT, Δ*cmaA* bacterial suspension (5 × 10^5^ CFU/ml), or water (mock) were infiltrated into three-week old cabbage and oat third leaf. Eight leaf discs (3.5 mm-diameter) from two cabbage leaves and about 3.5 cm^2^ from two oat leaves were collected 0 (untreated), 24 hpi, and 48 hpi, the weight was measured, and samples were frozen in liquid nitrogen and stored at −80°C. Samples were extracted with 300 μl of 80 % methanol.

SA was measured by using the multiple-reaction monitoring (MRM) mode on the LC-ESI-MS/MS (LCMS-8045; SHIMADZU, Kyoto, Japan) under the following conditions: capillary voltage, 4.5 kV; desolvation line, 250°C; heat block, 400°C; nebulizer nitrogen gas 3 L/min; drying gas, 10 L/min. Ion source polarity was set in the negative ion mode. The separation was performed with the LC system equipped with a 150 × 2.1 mm ACQUITY UPLC BEH Shield RP18 Column (Waters, Ireland) with a particle and pore size of 1.7 μm and 130Å, respectively. The initial mobile phase was solvent A: solvent B = 95:5 (solvent A, 0.5 mM ammonium formate (pH 3.5); solvent B, 0.5 mM ammonium formate (pH 3.5) in 90% acetonitrile). The solvent B concentration was increased to 100% for 8 min and then maintained at that ratio for another 2 min. The column was re-equilibrated for 3 min. The flow rate of 0.4 mL min^−1^ and the column temperature at 40°C were maintained throughout the analysis. The MRM-transition m/z 137→93 was used as precursor and product ions, respectively. The dwell time, Q1 pre bias, collision energy and Q3 pre bias were set at 100 msec, 15 V, 17 eV, 15 V, respectively.

### COR quantification by HPLC

*Pcal* WT and Δ*cmaA* bacteria were cultured in HS medium optimized for coronatine production (HSC; [82]) for 5 days. Culture supernatant was obtained by centrifugation (12,000 × g for 5 min). The culture supernatant was analyzed by HPLC with a Shimadzu LC20A system equipped with a Symmetry C8 column (4.6 × 250 mm; Waters Corporation, MA, USA) under the following conditions: column temperature, 40°C; isocratic elution; mobile-phase composition, 0.05% trifluoroacetic acid (TFA)/ acetonitrile (4:6, v/v); flow rate, 1 ml/min; and detection, 208 nm. COR in the culture supernatant was identified, as compared with authentic COR (Sigma) as the standard.

## Acknowledgments

We thank Dr. Christina Baker for editing the manuscript. *Pcal* was kindly given from the Nagano vegetable and ornamental crops experiment station, Nagano, Japan. This work was supported, in part, by JSPS KAKENHI Grant Number 19K0645, by Sustainable Food Security Research Project in the form of an operational grant from the National University Corporation, and by JST ERATO NOMURA Microbial Community Control Project, JST, Japan.

## Supporting information

**Supplementary Figure 1. HPLC-based quantification of COR production by *Pcal* WT and Δ*cmaA*.** *Pcal* WT and Δ*cmaA* bacteria were cultured in HS medium optimized for coronatine production (HSC; [81]) medium for 5 days. Culture supernatant was obtained by centrifugation, and analyzed by HPLC with a Shimadzu LC20A system equipped with a Symmetry C8 column under the following conditions: column temperature, 40°C; isocratic elution; mobile-phase composition, 0.05% trifluoroacetic acid (TFA)/ acetonitrile (4:6, v/v); flow rate, 1 ml/min; and detection, 208 nm. COR was identified in the culture supernatant, as compared with authentic COR as the standard. Red, green, and black chromatograms show authentic COR, *Pcal* WT, and *Pcal* Δ*cmaA*, respectively. Arrow indicates COR retention time.

**Supplementary Figure 2. COR reopens stomata on oat.** Stomatal opening rate of oat 1 h and 4 h after inoculation with *Pcal* WT and Δ*cmaA*. In all bar graphs, vertical bars indicate the standard error for three technical replicates.

**Supplementary Figure 3. COR contributes to cause disease on cabbage and oat. (A) and (B)** Disease symptoms on cabbage (A) and oat (B) 5 days after syringe-inoculation with *Pcal* WT and Δ*cmaA*. Plants were inoculated with 5 × 10^2^, 5 × 10^3^, 5 × 10^4^, and 5 × 10^5^ CFU/ml of *Pcal* WT and Δ*cmaA*. Scale bar shows 2 cm.

## References

1. Xin XF, Kvitko B, He SY. *Pseudomonas syringae*: what it takes to be a pathogen. Nat Rev Microbiol. 2018; 16: 316–328. doi: 10.1038/nrmicro.

2. Xin XF, He SY. *Pseudomonas syringae* pv. *tomato* DC3000: a model pathogen for probing disease susceptibility and hormone signaling in plants. Annu Rev Phytopathol. 2013; 51: 473–498. doi: 10.1146/annurev-phyto-082712-102321.

3. Bender CL, Palmer DA, Peñaloza-Vázquez A, Rangaswamy V, Ullrich M. Biosynthesis and regulation of coronatine, a non-host-specific phytotoxin produced by *Pseudomonas syringae*. Subcell Biochem. 1998; 29: 321–341. doi: 10.1007/978-1-4899-1707-2_10.

4. Buell CR, Joardar V, Lindeberg M, Selengut J, Paulsen IT, Gwinn ML, et al. The complete genome sequence of the Arabidopsis and tomato pathogen *Pseudomonas syringae* pv. *tomato* DC3000. Proc Natl Acad Sci U S A. 2003; 100: 10181–10186. doi: 10.1073/pnas.1731982100.

5. Collmer A, Lindeberg M, Petnicki-Ocwieja T, Schneider DJ, Alfano JR. Genomic mining type III secretion system effectors in *Pseudomonas syringae* yields new picks for all TTSS prospectors. Trends Microbiol. 2002; 10: 462–469. doi: 10.1016/s0966-842x(02)02451-4.

6. Budde IP, Ullrich MS. Interactions of *Pseudomonas syringae* pv. *glycinea* with host and nonhost plants in relation to temperature and phytotoxin synthesis. Mol Plant Microbe Interact. 2000; 13: 951–961. doi: 10.1094/MPMI.2000.13.9.951.

7. Mittal S, Davis KR. Role of the phytotoxin coronatine in the infection of *Arabidopsis thaliana* by *Pseudomonas syringae* pv. *tomato*. Mol Plant Microbe Interact. 1995; 8: 165–171. doi: 10.1094/mpmi-8-0165.

8. Elizabeth SV, Bender CL. The phytotoxin coronatine from *Pseudomonas syringae* pv. *tomato* DC3000 functions as a virulence factor and influences defence pathways in edible brassicas. Mol Plant Pathol. 2007; 8: 83–92. doi: 10.1111/j.1364-3703.2006.00372.x.

9. Sato M, Nishiyama K, Shirata A. Involvement of plasmid DNA in the productivity of coronatine by *Pseudomonas syringae* pv. *atropurpurea*. Ann Phytopathol Soc Japan. 1983; 49: 522–528. doi: 10.3186/jjphytopath.49.522

10. Ichihara A, Shiraishi K, Sato H, Sakamura S, Nishiyama K, Sakai R, et al. The structure of coronatine. J Am Chem Soc. 1977; 99: 636–637. doi: 10.1021/ja00444a067

11. Parry RJ, Mhaskar SV, Lin MT, Walker AE, Mafoti R. Investigations of the biosynthesis of the phytotoxin coronatine. Can J Chem. 1994; 72: 86–99. doi: 10.1139/v94-014.

12. Bender CL, Alarcón-Chaidez F, Gross DC. *Pseudomonas syringae* phytotoxins: mode of action, regulation, and biosynthesis by peptide and polyketide synthetases. Microbiol Mol Biol Rev. 1999; 63: 266–292.

13. Laurie-Berry N, Joardar V, Street IH, Kunkel BN. The *Arabidopsis thaliana* JASMONATE INSENSITIVE 1 gene is required for suppression of salicylic acid-dependent defenses during infection by *Pseudomonas syringae*. Mol Plant Microbe Interact. 2006; 19: 789–800. doi: 10.1094/MPMI-19-0789.

14. Feys B, Benedetti CE, Penfold CN, Turner JG. Arabidopsis mutants selected for resistance to the phytotoxin coronatine are male sterile, insensitive to methyl jasmonate, and resistant to a bacterial pathogen. Plant Cell. 1994; 6: 751–759. doi: 10.1105/tpc.6.5.751.

15. Weiler EW, Kutchan TM, Gorba T, Brodschelm W, Niesel U, Bublitz F. The *Pseudomonas* phytotoxin coronatine mimics octadecanoid signalling molecules of higher plants. FEBS Lett. 1994; 345: 9–13. doi: 10.1016/0014-5793(94)00411-0.

16. Melotto M, Underwood W, Koczan J, Nomura K, He SY. Plant stomata function in innate immunity against bacterial invasion. Cell. 2006; 126: 969–980. doi: 10.1016/j.cell.2006.06.054.

17. Mino Y, Matsushita Y, Sakai R. Effect of coronatine on stomatal opening in leaves of broadbean and italian ryegrass. Ann Phytopathol Soc Japan. 1987; 53: 53–55. doi: 10.3186/jjphytopath.53.53

18. Egoshi S, Takaoka Y, Saito H, et al. Dual function of coronatine as a bacterial virulence factor against plants: possible COI1–JAZ-independent role. RSC Adv. 2016; 6: 19404–19412. doi: 10.1039/C5RA20676F

19. Panchal S, Roy D, Chitrakar R, Price L, Breitbach ZS, Armstrong DW, et al. Coronatine facilitates *Pseudomonas syringae* infection of Arabidopsis leaves at night. Front Plant Sci. 2016; 7: 880. doi: 10.3389/fpls.2016.00880.

20. Toum L, Torres PS, Gallego SM, Benavídes MP, Vojnov AA, Gudesblat GE. Coronatine inhibits stomatal closure through guard cell-specific inhibition of NADPH oxidase-dependent ROS production. Front Plant Sci. 2016; 7: 1851. doi: 10.3389/fpls.2016.01851.

21. Dong X. SA, JA, ethylene, and disease resistance in plants. Curr Opin Plant Biol. 1998; 1: 316–323. doi: 10.1016/1369-5266(88)80053-0.

22. Glazebrook J. Contrasting mechanisms of defense against biotrophic and necrotrophic pathogens. Annu Rev Phytopathol. 2005; 43: 205–227. doi: 10.1146/annurev.phyto.43.040204.135923.

23. Thomma BP, Penninckx IA, Broekaert WF, Cammue BP. The complexity of disease signaling in Arabidopsis. Curr Opin Immunol. 2001; 13: 63–68. doi: 10.1016/s0952-7915(00)00183-7.

24. Kunkel BN, Brooks DM. Cross talk between signaling pathways in pathogen defense. Curr Opin Plant Biol. 2002; 5: 325–331. doi: 10.1016/s1369-5266(02)00275-3.

25. Brooks DM, Bender CL, Kunkel BN. The *Pseudomonas syringae* phytotoxin coronatine promotes virulence by overcoming salicylic acid-dependent defences in *Arabidopsis thaliana*. Mol Plant Pathol. 2005; 6: 629–639. doi: 10.1111/j.1364-3703.2005.00311.x.

26. Cui J, Bahrami AK, Pringle EG, Hernandez-Guzman G, Bender CL, Pierce NE, et al. *Pseudomonas syringae* manipulates systemic plant defenses against pathogens and herbivores. Proc Natl Acad Sci U S A. 2005; 102: 1791–1796. doi: 10.1073/pnas.0409450102.

27. Uppalapati SR, Ishiga Y, Wangdi T, Kunkel BN, Anand A, Mysore KS, et al. The phytotoxin coronatine contributes to pathogen fitness and is required for suppression of salicylic acid accumulation in tomato inoculated with *Pseudomonas syringae* pv. *tomato* DC3000. Mol Plant Microbe Interact. 2007; 20: 955–965. doi: 10.1094/MPMI-20-8-0955.

28. Ishiga Y, Ishiga T, Wangdi T, Mysore KS, Uppalapati SR. NTRC and chloroplast-generated reactive oxygen species regulate *Pseudomonas syringae* pv. *tomato* disease development in tomato and Arabidopsis. Mol Plant Microbe Interact. 2012; 25: 294–306. doi: 10.1094/MPMI-05-11-0130.

29. Zheng XY, Spivey NW, Zeng W, Liu PP, Fu ZQ, Klessig DF, et al. Coronatine promotes *Pseudomonas syringae* virulence in plants by activating a signaling cascade that inhibits salicylic acid accumulation. Cell Host Microbe. 2012; 11: 587–596. doi: 10.1016/j.chom.2012.04.014.

30. Bender CL, Liyanage H, Palmer D, Ullrich M, Young S, Mitchell R. Characterization of the genes controlling the biosynthesis of the polyketide phytotoxin coronatine including conjugation between coronafacic and coronamic acid. Gene. 1993; 133: 31–38. doi: 10.1016/0378-1119(93)90221-n.

31. Brooks DM, Hernández-Guzmán G, Kloek AP, Alarcón-Chaidez F, Sreedharan A, Rangaswamy V, et al. Identification and characterization of a well-defined series of coronatine biosynthetic mutants of *Pseudomonas syringae* pv. *tomato* DC3000. Mol Plant Microbe Interact. 2004; 17: 162–174. doi: 10.1094/MPMI.2004.17.2.162.

32. Sreedharan A, Penaloza-Vazquez A, Kunkel BN, Bender CL. CorR regulates multiple components of virulence in *Pseudomonas syringae* pv. *tomato* DC3000. Mol Plant Microbe Interact. 2006; 19: 768–779. doi: 10.1094/MPMI-19-0768.

33. Peñaloza-Vázquez A, Bender CL. Characterization of CorR, a transcriptional activator which is required for biosynthesis of the phytotoxin coronatine. J Bacteriol. 1998; 180:6252–6259. doi: 10.1128/JB.180.23.6252-6259.1998

34. Xiao Y, Heu S, Yi J, Lu Y, Hutcheson SW. Identification of a putative alternate sigma factor and characterization of a multicomponent regulatory cascade controlling the expression of *Pseudomonas syringae* pv. *syringae* Pss61 *hrp* and *hrmA* genes. J Bacteriol. 1994; 176: 1025–1036. doi: 10.1128/jb.176.4.1025-1036.1994.

35. Peñaloza-Vázquez A, Preston GM, Collmer A, Bender CL. Regulatory interactions between the Hrp type III protein secretion system and coronatine biosynthesis in *Pseudomonas syringae* pv. *tomato* DC3000. Microbiology. 2000; 146: 2447–2456. doi: 10.1099/00221287-146-10-2447.

36. Bull CT, Manceau C, Lydon J, Kong H, Vinatzer BA, Fischer-Le Saux M. *Pseudomonas cannabina* pv. *cannabina* pv. nov., and *Pseudomonas cannabina* pv. *alisalensis* (Cintas Koike and Bull, 2000) comb. nov., are members of the emended species *Pseudomonas cannabina* (ex Sutic & Dowson 1959) Gardan, Shafik, Belouin, Brosch, Grimont & Grimont 1999. Syst Appl Microbiol. 2010; 33: 105–115. doi: 10.1016/j.syapm.2010.02.001.

37. Cintas NA, Koike ST, Bull CT. A new pathovar, *Pseudomonas syringae* pv. *alisalensis* pv. nov., proposed for the causal agent of bacterial blight of broccoli and broccoli raab. Plant Dis. 2002; 86: 992–998. doi: 10.1094/PDIS.2002.86.9.992.

38. Sarris PF, Trantas EA, Baltrus DA, Bull CT, Wechter WP, Yan S, et al. Comparative genomics of multiple strains of *Pseudomonas cannabina* pv. *alisalensis*, a potential model pathogen of both monocots and dicots. PLoS One. 2013; 8: e59366. doi: 10.1371/journal.pone.0059366.

39. Takahashi F, Ochiai M, Ikeda K, Takikawa Y. Streptomycin and copper resistance in *Pseudomonas cannabina* pv. *alisalensis* (abstract in Japanese). Jpn J Phytopathol. 2013. p. 35.

40. Takikawa Y, Takahashi F. Bacterial leaf spot and blight of crucifer plants (Brassicaceae) caused by *Pseudomonas syringae* pv. m*aculicola* and *P.* c*annabina* pv. *alisalensis*. J Gen Plant Pathol. 2014; 80: 466–474. doi: 10.1007/s10327-014-0540-4.

41. Ishiyama Y, Yamagishi N, Ogiso H, Fujinaga M, Takahashi F, Takikawa Y. Bacterial brown spot on *Avena storigosa* Schereb. caused by *Pseudomonas syringae* pv. *alisalensis*. J Gen Plant Pathol. 2013; 79: 155–157. doi:10.1007/s10327-013-0427-9.

42. Ishiga T, Iida Y, Sakata N, Ugajin T, Hirata T, Taniguchi S, et al. Acibenzolar-S-methyl activates stomatal-based defense against bacterial pathogen in cabbage. J Gen Plant Pathol 2020; 86, 48–54. doi: 10.1007/s10327-019-00883-5.

43. Ishiga T, Iida Y, Sakata N, Ugajin T, Hirata T, Taniguchi S, et al. Acibenzolar-S-methyl and probenazole control bacterial blight disease of cabbage by activating stomatal-based defense in different timing manners. J Gen Plant Pathol. 2020. Forthcoming

44. Sakata N, Ishiga T, Taniguchi S, Ishiga Y. Acibenzolar-S-methyl activates stomatal-based defense systemically in Japanese radish by inducing peroxidase-dependent reactive oxygen species production. bioRxiv. 2020:2020.05.24.113878. doi: 10.1101/2020.05.24.113878.

45. Sakata N, Ishiga T, Saito H, Nguyen VT, Ishiga Y. reveals *Pseudomonas cannabina* pv. *alisalensis* optimizes its virulence factors for pathogenicity on different hosts. PeerJ. 2019; 7: e7698. doi: 10.7717/peerj.7698.

46. Evidence for a transmissible factor that causes rapid stomatal closure in soybean at sites adjacent to and remote from hypersensitive cell death induced by *Phytophthora sojae*. Physiol Mol Plant P. 1999; 55: 197–203. doi: 10.1006/pmpp.1999.0220.

47. Gudesblat GE, Torres PS, Vojnov AA. Stomata and pathogens: Warfare at the gates. Plant Signal Behav. 2009; 4: 1114–1116. doi: 10.4161/psb.4.12.10062.

48. Zhang H, Dong S, Wang M, Wang W, Song W, Dou X, et al. The role of vacuolar processing enzyme (VPE) from *Nicotiana benthamiana* in the elicitor-triggered hypersensitive response and stomatal closure. J Exp Bot. 2010; 61: 3799–3812. doi: 10.1093/jxb/erq189.

49. Roy D, Panchal S, Rosa BA, Melotto M. *Escherichia coli* O157:H7 induces stronger plant immunity than *Salmonella enterica* Typhimurium SL1344. Phytopathology. 2013; 103: 326–332. doi: 10.1094/PHYTO-09-12-0230-FI.

50. Arnaud D, Hwang I. A sophisticated network of signaling pathways regulates stomatal defenses to bacterial pathogens. Mol Plant. 2015; 8: 566–581. doi: 10.1016/j.molp.2014.10.012.

51. Sawinski K, Mersmann S, Robatzek S, Böhmer M. Guarding the green: pathways to stomatal immunity. Mol Plant Microbe Interact. 2013; 26: 626–632. doi: 10.1094/MPMI-12-12-0288-CR.

52. Zhang D, Tian C, Yin K, Wang W, Qiu JL. Postinvasive bacterial resistance conferred by open stomata in rice. Mol Plant Microbe Interact. 2019; 32: 255–266. doi: 10.1094/MPMI-06-18-0162-R.

53. Helmann TC, Deutschbauer A, Lindow S. Distinctiveness of genes contributing to growth of *Pseudomonas syringae* in diverse host plant species. bioRxiv. 2020:2020.07.28.216440. doi: 10.1101/2020.07.28.216440.

54. Uppalapati SR, Ayoubi P, Weng H, Palmer DA, Mitchell RE, Jones W, et al. The phytotoxin coronatine and methyl jasmonate impact multiple phytohormone pathways in tomato. Plant J. 2005; 42: 201–217. doi: 10.1111/j.1365-313X.2005.02366.x..

55. Kloek AP, Verbsky ML, Sharma SB, Schoelz JE, Vogel J, Klessig DF, et al. Resistance to *Pseudomonas syringae* conferred by an *Arabidopsis thaliana* coronatine-insensitive (*coi1*) mutation occurs through two distinct mechanisms. Plant J. 2001; 26: 509–522. doi: 10.1046/j.1365-313x.2001.01050.

56. De Vleesschauwer D, Xu J, Höfte M. Making sense of hormone-mediated defense networking: from rice to Arabidopsis. Front Plant Sci. 2014; 5: 611. doi: 10.3389/fpls.2014.00611.

57. Silverman P, Seskar M, Kanter D, Schweizer P, Metraux JP, Raskin I. Salicylic acid in rice (biosynthesis, conjugation, and possible role). Plant Physiol. 1995; 108: 633–639. doi: 10.1104/pp.108.2.633.

58. Malamy J, Klessig DF. Salicylic acid and plant disease resistance. Plant J. 1992; 2: 643–654. doi: 10.1111/j.1365-313X.1992.tb00133.x.

59. Lee MW, Qi M, Yang Y. A novel jasmonic acid-inducible rice myb gene associates with fungal infection and host cell death. Mol Plant Microbe Interact. 2001; 14: 527–535. doi: 10.1094/MPMI.2001.14.4.527.

60. Ding L, Yang G, Yang R, Cao J, Zhou Y. Investigating interactions of salicylic acid and jasmonic acid signaling pathways in monocots wheat. Physiol Mol Plant P. 2016; 93: 67–74. doi: 10.1016/j.pmpp.2016.01.002.

61. Iwai T, Seo S, Mitsuhara I, Ohashi Y. Probenazole-induced accumulation of salicylic acid confers resistance to *Magnaporthe grisea* in adult rice plants. Plant Cell Physiol. 2007; 48: 915–924. doi: 10.1093/pcp/pcm062.

62. Geng X, Cheng J, Gangadharan A, Mackey D. The coronatine toxin of *Pseudomonas syringae* is a multifunctional suppressor of Arabidopsis defense. Plant Cell. 2012; 24: 4763–4774. doi: 10.1105/tpc.112.105312.

63. Thilmony R, Underwood W, He SY. Genome-wide transcriptional analysis of the *Arabidopsis thaliana* interaction with the plant pathogen *Pseudomonas syringae* pv. *tomato* DC3000 and the human pathogen *Escherichia coli* O157:H7. Plant J. 2006; 46: 34–53. doi: 10.1111/j.1365-313X.2006.02725.x.

64. Klein AP, Sattely ES. Biosynthesis of cabbage phytoalexins from indole glucosinolate. Proc Natl Acad Sci U S A. 2017; 114: 1910–1915. doi: 10.1073/pnas.1615625114.

65. Pedras MSC, Abdoli A, Sarma-Mamillapalle VK. Inhibitors of the detoxifying enzyme of the phytoalexin brassinin based on quinoline and isoquinoline scaffolds. Molecules. 2017; 22: 1345. doi: 10.3390/molecules22081345.

66. Wang W, Yang J, Zhang J, Liu YX, Tian C, Qu B, et al. An Arabidopsis secondary metabolite directly targets expression of the bacterial type III secretion system to inhibit bacterial virulence. Cell Host Microbe. 2020; 27: 601–613.e7. doi: 10.1016/j.chom.2020.03.004.

67. Collins FW. Oat phenolics: avenanthramides, novel substituted N-cinnamoylanthranilate alkaloids from oat groats and hulls. J Agric Food Chem. 1989; 37: 60–66. doi: 10.1021/jf00085a015.

68. Wise ML. Effect of chemical systemic acquired resistance elicitors on avenanthramide biosynthesis in oat (*Avena sativa*). J Agric Food Chem. 2011; 59: 7028–7038. doi: 10.1021/jf2008869.

69. Matsukawa T, Ishihara A, Iwamura H. Induction of anthranilate synthase activity by elicitors in oats. Z Naturforsch C J Biosci. 2002; 57: 121–128. doi: 10.1515/znc-2002-1-221.

70. Osbourn A, Goss RJ, Field RA. The saponins: polar isoprenoids with important and diverse biological activities. Nat Prod Rep. 2011; 28: 1261–1268. doi: 10.1039/c1np00015b.

71. Faizal A, Geelen D. Saponins and their role in biological processes in plants. Phytochemistry reviews. 2013; 12: 877–893. doi: 10.1007/s11101-013-9322-4.

72. King E, Ward M, Raney D. Two simple media for the demonstration of pyocyanin and fluorescin. J Lab Clin Med. 1954; 44: 301–307.

73. Sambrook J, Fritsch E, Maniatis T. Molecular cloning: a laboratory manual, 2nd edition.: Cold Spring Harbor Laboratory, Cold Spring Harbor, New York; 1989.

74. Wang K, Kang L, Anand A, Lazarovits G, Mysore KS. Monitoring in planta bacterial infection at both cellular and whole-plant levels using the green fluorescent protein variant GFPuv. New Phytol. 2007; 174: 212–223. doi: 10.1111/j.1469-8137.2007.01999.x.

75. Ishiga T, Ishiga Y, Betsuyaku S, Nomura N. AlgU contributes to the virulence of *Pseudomonas syringae* pv. *tomato* DC3000 by regulating phytotoxin coronatine production. J Gen Plant Pathol. 2018; 84: 189–201. doi: 10.1007/s10327-018-0775-6.

76. Schäfer A, Tauch A, Jäger W, Kalinowski J, Thierbach G, Pühler A. Small mobilizable multi-purpose cloning vectors derived from the *Escherichia coli* plasmids pK18 and pK19: selection of defined deletions in the chromosome of *Corynebacterium glutamicum*. Gene. 1994; 145: 69–73. doi: 10.1016/0378-1119(94)90324-7.

77. Shimizu R, Taguchi F, Marutani M, Mukaihara T, Inagaki Y, Toyoda K, et al. The DeltafliD mutant of *Pseudomonas syringae* pv. *tabaci*, which secretes flagellin monomers, induces a strong hypersensitive reaction (HR) in non-host tomato cells. Mol Genet Genomics. 2003; 269: 21–30. doi: 10.1007/s00438-003-0817-3.

78. Keane PJ, Kerr A, New PB. Crown gall of stone fruit. II. Identification and nomenclature of *Agrobacterium* isolates. Aust J Biol Sci. 1979; 23: 585–595.

79. Ishiga Y, Ichinose Y. *Pseudomonas syringae* pv. *tomato* OxyR is required for virulence in tomato and Arabidopsis. Mol Plant Microbe In. 2016; 29: 119–131. doi: 10.1094/MPMI-09-15-0204-R.

80. Ishiga Y, Uppalapati SR, Ishiga T, Elavarthi S, Martin B, Bender CL. The phytotoxin coronatine induces light-dependent reactive oxygen species in tomato seedlings. New Phytol. 2009; 181: 147–160. doi: 10.1111/j.1469-8137.2008.02639.x.

81. Chitrakar R, Melotto M. Assessing stomatal response to live bacterial cells using whole leaf imaging. J Vis Exp. 2010; 44: e2185. doi: 10.3791/2185.

82. Palmer DA, Bender CL. Effects of environmental and nutritional factors on production of the polyketide phytotoxin coronatine by *Pseudomonas syringae* pv. *glycinea*. Appl Environ Microbiol. 1993; 59: 1619–1626. doi: 10.1128/AEM.59.5.1619-1626.1993.

