## Supplementary figures and images for "Coronatine contributes *Pseudomonas cannabina* pv. *alisalensis* virulence by overcoming both stomatal and apoplastic defenses in dicot and monocot plants"

### Supplemental Figures

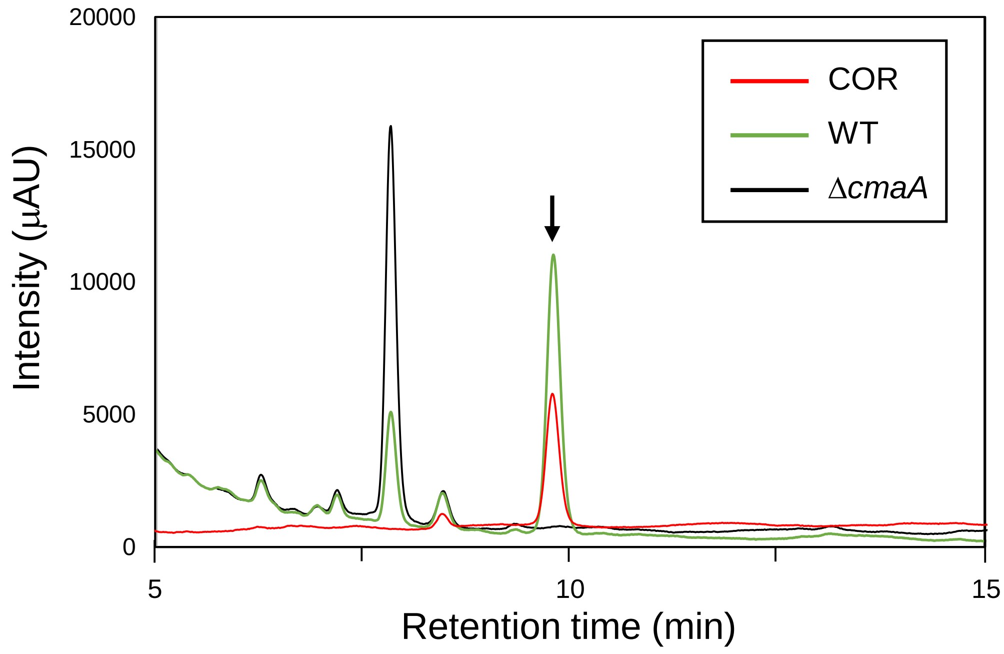

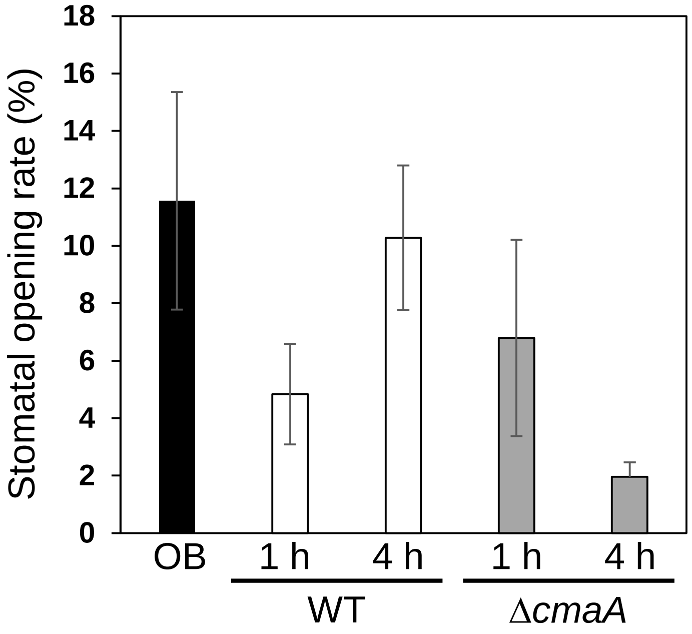

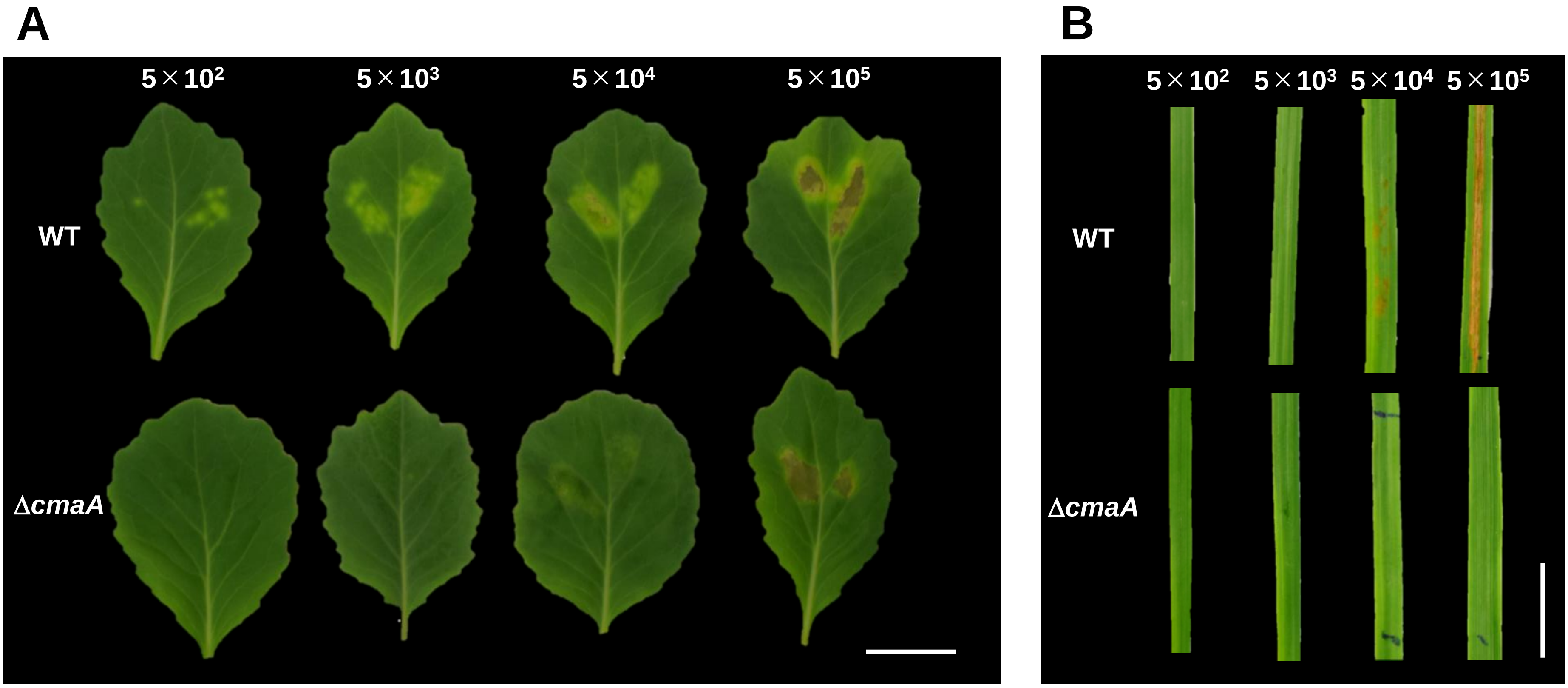
