## Supplemental Tables for "Coronatine contributes *Pseudomonas cannabina* pv. *alisalensis* virulence by overcoming both stomatal and apoplastic defenses in dicot and monocot plants"

**Supplementary Table S1** Bacterial strains and plasmids used in this study

| Bacterial strain or plasmid | Relevant characteristics | Reference or source |
|---|---|---|
| *E. coli* strain | | |
| DH5α | *F*$^{-}$ *λ*$^{-}$ *ϕ80dLac*ZΔM15 Δ(*lacZYA-argF*)U169 *recA*1 *endA*1 *hsdR*17 (rK$^{-}$ mK$^{+}$) *supE*44 *thi-1gyrA relA*1 | Takara, Kyoto, Japan |
| S17-1 | *thi pro hsdR*$^{-}$ *hsdM*$^{+}$ *recA* [chr::RP4-2-Tc::Mu-Km::Tn7] | Schafer et al. (1994) |
| *P. cannabina* pv. *alisalensis* (*Pcal*) | | |
| Isolate KB211 | Wild type, Rif $^{r}$ | Nagano vegetable and ornamental crops experiment station |
| KB211 tdTomato | *Pcal* KB211 containing pDSK-tdTomato, Rif $^{r}$, Km$^{r}$ | This work |
| KB211 Δ*cmaA* | *Pcal* KB211 Δ*cmaA*, Rif $^{r}$ | This work |
| Plasmid | | |
| pGEM-T Easy | Cloning vector, Amp$^{r}$ | Promega, Tokyo, japan |
| pK18*mobsacB* | Small mobilizable vector, Km$^{r}$, sucrose sensitive (*sacB*) | Schafer et al. (1994) |
| pDSK-*tdTomato* | The vector containing constitutive *psbA* promoter and *tdTomato* gene inserted into pDSK519, Km$^{r}$ | This work |

Amp$^{r}$ ampicillin resistance, Km$^{r}$ kanamycin resistance, Rif $^{r}$ rifampicin resistance

**Supplementary Table S2** Primer sets used in this study

| Gene | Primer name | Primer usage | Primer sequence | Reference |
|---|---|---|---|---|
| *cmaA* | cmaA-P1 | Deletion mutagenesis of *cmaA* | AGTGATTGCGGATCAGGTTAAG | This study |
| *cmaA* | cmaA-P2 | Deletion mutagenesis of *cmaA* | TCGCTGTGGCACTGATTT | This study |
| *cmaA* | cmaA-P3 | Deletion mutagenesis of *cmaA* | CGGGATCCTGTGAATGGTAGGAGGTCAT | This study |
| *cmaA* | cmaA-P4 | Deletion mutagenesis of *cmaA* | CGGGATCCGCCAACATGGAAATGACTGA | This study |
| *cmaA* | cmaA-FW | Expression analysis on *Pcal* | AAAGCCTACCGCCGATTT | This study |
| *cmaA* | cmaA-RV | Expression analysis on *Pcal* | CGTCTGGAGCTGTTGATAAGT | This study |
| *cfl* | cfl-FW | Expression analysis on *Pcal* | GAACTGGTGGCGTTGTACTAT | This study |
| *cfl* | cfl-RV | Expression analysis on *Pcal* | GTGGAGCAGATGCTCAATTTC | This study |
| *corR* | corR-FW | Expression analysis on *Pcal* | GGATCGAACGCTGGCAGATA | This study |
| *corR* | corR-RV | Expression analysis on *Pcal* | GTCCTGCTCATGAGTCGCTT | This study |
| *hrpL* | hrpL-FW | Expression analysis on *Pcal* | CATGCCAGTAAACCGCAAAC | This study |
| *hrpL* | hrpL-RV | Expression analysis on *Pcal* | GTCTTCCCAGCTTTCCTGATAA | This study |
| *oprF* | oprF-FW | Expression analysis on *Pcal* | GGCTTGGCCATTGGTACTAT | This study |
| *oprF* | oprF-RV | Expression analysis on *Pcal* | GCGCTGTCGTAATACTCTTTCT | This study |
| *recA* | recA-FW | Expression analysis on *Pcal* | TCTCTACGGCAAGGGTATCT | This study |
| *recA* | recA-RV | Expression analysis on *Pcal* | GCTTTACCCTGACCGATCTT | This study |
| *BoPR1* | BoPR1-FW | Expression analysis on cabbage | GGTCAACGAGAAGGCTAACTATAA | Ishiga et al., 2020 |
| *BoPR1* | BoPR1-RV | Expression analysis on cabbage | GCTTTGCCACATCCAATTCTC | Ishiga et al., 2020 |
| *BoPR2* | BoPR2-FW | Expression analysis on cabbage | GAAGAGTGGAACTCCGAGAAAG | Ishiga et al., 2020 |
| *BoPR2* | BoPR2-RV | Expression analysis on cabbage | AGGCTGTTGACTAGGAAGAAAC | Ishiga et al., 2020 |
| *BoUBQ1* | BoUBQ1-FW | Expression analysis on cabbage | GTCAAGGCCAAGATCCAAGA | Ishiga et al., 2020 |
| *BoUBQ1* | BoUBQ1-RV | Expression analysis on cabbage | GGATGTTGTAGTCAGCCAGAG | Ishiga et al., 2020 |
| *AsPR1a* | AsPR1a-FW | Expression analysis on oat | AGCTACGCCGAGCAGAG | This study |
| *AsPR1a* | AsPR1a-RV | Expression analysis on oat | ATGGTCGTAGTACTGCTTCTCC | This study |
| *AsPR1b* | AsPR1b-FW | Expression analysis on oat | CTCGGCTCAAGACTACCTTTC | This study |
| *AsPR1b* | AsPR1b-RV | Expression analysis on oat | TAGCTCTCCGCATACGACT | This study |
| *AsPR2* | AsPR2-FW | Expression analysis on oat | TTCAGTGGCATCAAGGTCTC | This study |
| *AsPR2* | AsPR2-RV | Expression analysis on oat | CGGTGCTCTCCAGAAACTC | This study |
| *AsAct* | AsAct-FW | Expression analysis on oat | CATGAAGTGCGACGTGGATA | This study |
| *AsAct* | AsAct-RV | Expression analysis on oat | CGGTGATCTCCTTGCTCATAC | This study |

BamHI digestion sites are underlined.
